# Physical constraints and biological regulations underlie universal osmoresponses

**DOI:** 10.1101/2024.07.02.601668

**Authors:** Yiyang Ye, Qirun Wang, Jie Lin

## Abstract

Microorganisms constantly transition between environments with dramatically different external osmolarities. However, theories of microbial osmoresponse integrating physical constraints and biological regulations are lacking. Here, we propose such a theory, utilizing the separation of timescales for passive responses and active regulations. We demonstrate that regulations of osmolyte production and cell-wall synthesis assist cells in coping with intracellular crowding effects and adapting to a broad range of external osmolarity. Furthermore, we predict a threshold value above which cells cannot grow, ubiquitous across bacteria and yeast. Intriguingly, the theory predicts a dramatic speedup of cell growth after an abrupt decrease in external osmolarity due to cell-wall synthesis regulation. Our theory rationalizes the unusually fast growth observed in fission yeast after an oscillatory osmotic perturbation, and the predicted growth rate peaks match quantitatively with experimental measurements. Our study reveals the physical basis of osmoresponse, yielding far-reaching implications for microbial physiology.

---

Microbes constantly transition between environments with dramatically different osmolarities, a hallmark of microbial life [1–4]. One of the most essential features of walled microbial cells is the turgor pressure — the elastic stress stretching the cell wall due to osmotic imbalance. Upon a hypoosmotic shock (i.e., a sudden decrease of the external osmolarity), the turgor pressure increases immediately due to the sudden water influx. To relax the turgor pressure, the cell up-regulates the cell-wall synthesis rate, adds more materials to the peptidoglycan network, and eventually adapts to the lower external osmolarity [5]. Upon a hyperosmotic shock (i.e., a sudden increase of the external osmolarity), the cell volume of a microbial cell shrinks within milliseconds due to water efflux, leading to a decreased turgor pressure [6, 7]. To increase the internal osmotic pressure, microorganisms increase their intracellular solute pool by amassing osmolyte molecules (i.e., osmoregulation), e.g., through *de novo* synthesis [8]. The cell volume then restores progressively over time, and eventually, the cell adapts to the higher osmolarity. Intracellular crowding may act as a cell volume sensor to trigger osmoregulation [9–11]. Meanwhile, intracellular crowding due to volume reduction inevitably affects the cellular physiology globally, e.g., slowing down protein diffusion [12–18] and reducing the elongation speed of translating ribosomes [19, 20]. Despite extensive knowledge regarding the molecular details of osmotic response pathways [4], how intracellular crowding interferes with gene expression regulation and affects osmotic adaption remains an open question.

Interestingly, many features of microbial osmoresponses appear general across different organisms, suggesting a universal underlying mechanism. For example, it is widely observed that microbial cells can adapt to a broad range of external osmolarity, with the external osmotic pressure varying over an order of magnitude [19, 21–23]. Furthermore, the growth rate in the steady state decreases as the external osmolarity increases and a complete arrest of cell growth occurs above a critical osmolarity [19, 22–26]. Moreover, upon an osmotic shock, the growth rate usually does not approach the new steady-state value monotonically, e.g., an overshoot of growth rate often occurs upon a hypoosmotic shock [23], and a damped oscillation of growth rate can happen after a hyperosmotic shock [22]. In recent experiments of *Schizosaccharomyces pombe* [27], an oscillatory osmotic shock was applied to cells during which cell volume growth was dramatically slowed down while biomass was still actively produced. Surprisingly, a supergrowth phase happened after removing the oscillatory osmotic shock, during which cells grew much faster than the steady state before the shocks.

In this work, we unify all these phenomena by a theory capturing the essential elements of osmoresponses: physical constraints (e.g., the crowding effects and osmotic imbalance) and biological regulation, including osmoregulation (i.e., regulation of the osmolyte-producing protein) and cell-wall synthesis regulation. Our model assumes the following phenomenological rules: (1) the change in free water volume within the cell is driven by osmotic imbalance [7, 28], while the remaining volume changes in proportion to protein production; (2) osmoregulation influences the production of osmolyte-producing protein, governed by intracellular protein density [29]; (3) cell-wall synthesis is regulated through a feedback mechanism, wherein turgor pressure modulates the efficiency of cell-wall synthesis, enabling the cell to maintain a relatively stable turgor pressure; and (4) intracellular crowding slows down biochemical reactions as cytoplasmic density increases, with reactions ceasing entirely when protein density reaches a critical threshold. Upon a hyperosmotic shock, cell volume reduction due to water efflux increases the protein density, inducing the up-regulation of osmolyte-producing protein but slowing down the translation speed due to crowding. Upon a hypoosmotic shock, the dramatic water influx stretches the cell wall, and the increased turgor pressure induces cell-wall synthesis [5, 30, 31].

We remark that our model is coarse-grained, without including detailed molecular mechanisms, and is therefore applicable across diverse microbial species. Notably, the predicted steady-state growth rate as a function of internal osmotic pressure from our model aligns well with experimental data from diverse organisms. This alignment allows us to quantify the sensitivities of translation speed and regulation of osmolyte-producing protein in response to intracellular density. Additionally, we demonstrate that osmoregulation and cell-wall synthesis regulation enable cells to adapt to a wide range of external osmolarities and prevent plasmolysis. Our model also predicts a non-monotonic time dependence of growth rate and protein density as they approach steady-state values following a constant osmotic shock, in concert with experimental observations [22, 23]. Moreover, we show that a supergrowth phase can arise following a sudden decrease in external osmolarity, driven by cell-wall synthesis regulation, either through the direct application of a hypoosmotic shock or the withdrawal of an oscillatory stimulus. Remarkably, the predicted amplitudes of supergrowth (i.e., growth rate peaks) quantitatively agree with multiple independent experimental measurements.

In the following “Results” section, we begin by outlining the primary assumptions and equations of our model in the subsection “Model Description,” which includes four parts, each addressing one of the four phenomenological rules. Additional details can be found in Methods. We then proceed to the subsection “Steady states in constant environments”, where we employ our theoretical framework to analyze steady-state growth and examine how the growth rate varies with external osmolarity. In the “Transient dynamics after a constant osmotic shock” subsection, we investigate the time-dependent osmoresponse after a constant hyperosmotic and hypoosmotic shock. Finally, in “Comparison with experiments: supergrowth phenomena after osmotic oscillation”, we address the supergrowth phenomena observed in *S. pombe*, utilizing our model to elucidate these experimental observations.

## RESULTS

### Model Description

#### Cell growth

In the limit of an extreme hyperosmotic shock, the remaining cytoplasmic volume is comparable to the volume of expelled water [14, 21, 32]. Thus, the total cytoplasmic volume must be divided into a free volume and a bound volume [28, 33–36],

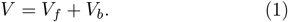

The free volume comes from the free water that is osmotically active, and the bound volume includes the bound water *V*_*bw*_ (i.e., water of macromolecular hydration) and the volume of dry mass *V*_*bd*_ (Figure 1A). Because the fraction of protein mass in the total dry mass is typically constant and the volume of bound water is proportional to the dry mass [21], the bound volume is proportional to the total protein mass *m*_*p*_ through *V*_*b*_ = *αm*_*p*_. Here, *α* is a constant, and its values for some model organisms are included in Table 1, and its detailed calculations from experimental data are in Section B of Supplementary Material.

**TABLE 1.**
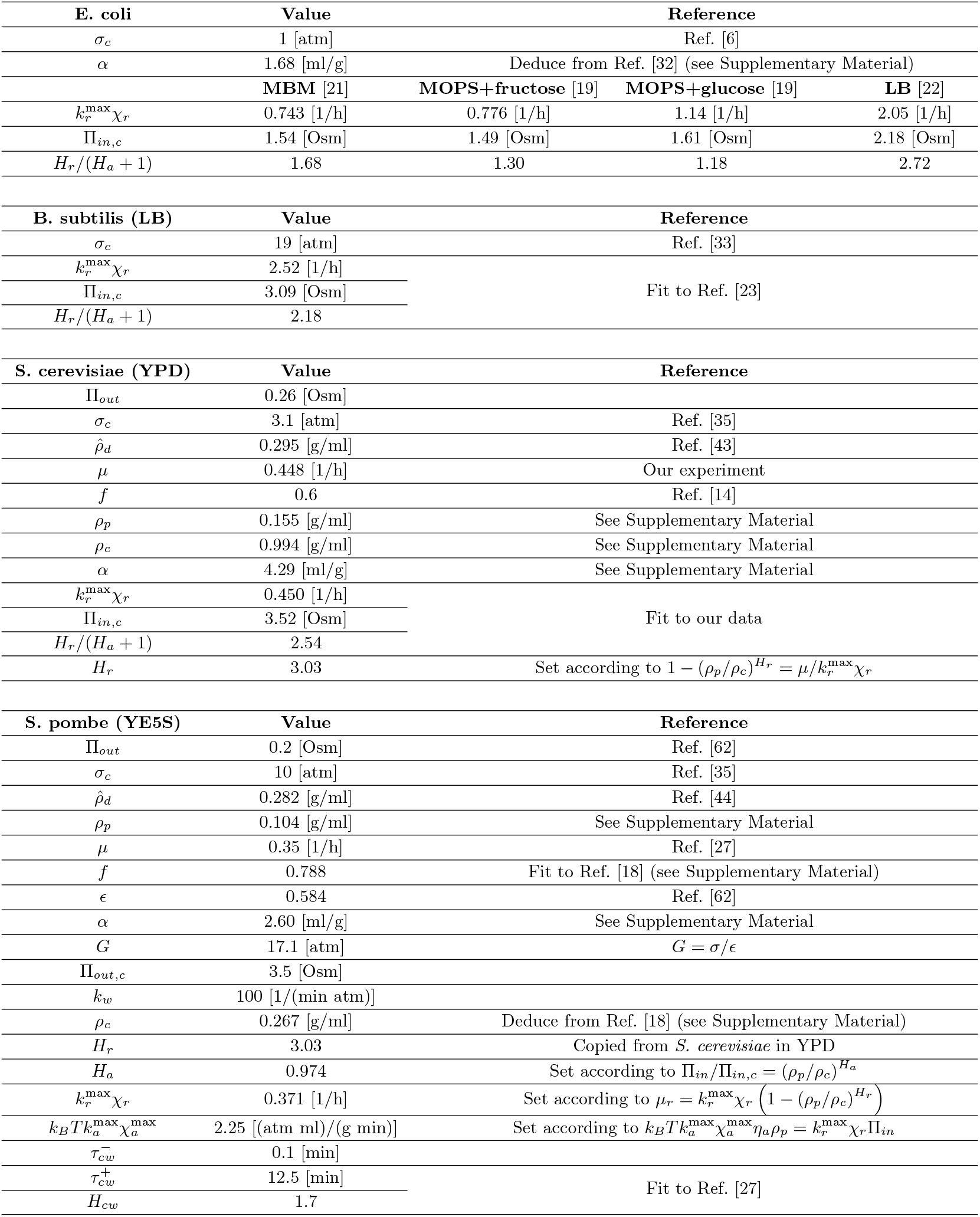
Model parameters for different species in their corresponding reference growth media.

**FIG. 1.**
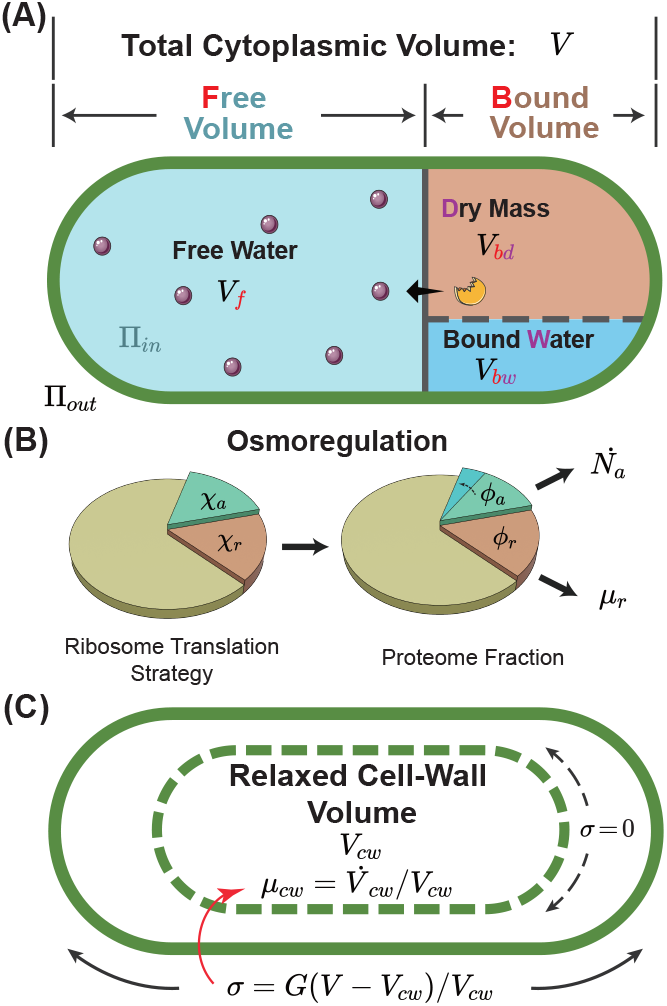
A schematic of the osmoresponse model. (A) The total cytoplasmic volume includes the free and bound volumes. The free volume sets the internal osmotic pressure Π_*in*_ = *k*_*B*_*TN*_*a*_*/V*_*f*_, where *V*_*f*_ is the free volume and *N*_*a*_ is the number of osmolyte molecules. The bound volume *V*_*b*_ comprises the dry mass *V*_*bd*_ and bound water *V*_*bw*_, i.e., *V*_*b*_ = *V*_*bd*_ +*V*_*bw*_, all proportional to the total protein mass. We model osmoregulation through the change of ribosome translation strategy. When the protein density increases, the fraction of ribosomes translating the osmolyte-producing protein *χ*_*a*_ is up-regulated, leading to the subsequent increase in the mass fraction of the osmolyte-producing protein *ϕ*_*a*_. The cell-wall synthesis process is controlled by the turgor pressure *σ*, which is proportional to the cell-wall strain *ϵ* = (*V* −*V*_*cw*_)*/V*_*cw*_ . Here, *V* is the cytoplasmic volume, and *V*_*cw*_ is the relaxed cell wall volume.

The free volume changes due to osmotic imbalance, and the growth rate of the free volume follows

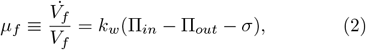

where Π_*in*_, Π_*out*_ are the internal (i.e., cytoplasmic) and external osmotic pressures, respectively [7]. Π_*in*_ is proportional to the concentration of osmolyte molecules in the free volume: Π_*in*_ = *k*_*B*_*TN*_*a*_*/V*_*f*_ where *N*_*a*_ is the number of osmolyte molecules, *k*_*B*_ is the Boltzmann constant, and *T* is the temperature. For simplicity, we assume that the production speed of osmolyte molecules is proportional to the mass of osmolyte-producing protein (Methods). Here, we have replaced the difference of the hydrostatic pressures across the cell membrane with the turgor pressure *σ*, assuming that mechanical equilibrium is always satisfied. *k*_*w*_ is the filtration coefficient quantifying the water permeability of the cell membrane [37].

The species of osmolytes involved in osmoregulation are diverse across different microorganisms and conditions; nevertheless, they are primarily small organic molecules [8, 38]. In this work, we simplify the problem by considering a single species of osmolyte that dominates the internal osmotic pressure, e.g., glycerol in *Saccharomyces cerevisiae* [39–41] and glycine betaine in *Escherichia coli* [3], with the production speed proportional to the mass of the osmolyte-producing protein (Figure 1A and Methods).

To model gene expression regulation, we introduce *χ*_*a*_ and *χ*_*r*_ as the fractions of ribosomes translating the osmolyte-producing protein and ribosomal proteins (Figure 1B and Methods). In steady states, *χ*_*a*_ and *χ*_*r*_ are equal to the mass fractions of osmolyte-producing protein and ribosomal proteins in the total proteome, *ϕ*_*a*_ = *m*_*p*,*a*_*/m*_*p*_ and *ϕ*_*r*_ = *m*_*p*,*r*_*/m*_*p*_, respectively [29, 42]. In this work, we assume that the dry-mass growth rate is proportional to the fraction of ribosomal proteins within the total proteome for simplicity, *µ*_*r*_ = *k*_*r*_*m*_*p*,*r*_*/m*_*p*_ = *k*_*r*_*ϕ*_*r*_. This assumption leverages the fact that ribosomes are responsible for producing all proteins. The proportionality coefficient *k*_*r*_ encapsulates the efficiency of ribosomal activity, being proportional to the elongation speed of the ribosome. We remark that *k*_*r*_ is influenced by the crowding effect, which we address later. The growth rate of the cytoplasmic volume, 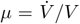 is a weighted average of the free-volume growth rate *µ*_*f*_ and the dry-mass growth rate *µ*_*r*_.

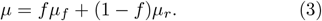

Here, *f* is the free volume fraction in the total cytoplasmic volume: *f* = *V*_*f*_ */V* . In this work, we refer to the growth rate as the growth rate of cytoplasmic volume *µ* unless otherwise mentioned.

#### Osmoregulation

Experiments found that the reduction of growth rate as the external osmolarity increases is dominated by the reduction of the translation speed *k*_*r*_ instead of the ribosomal fraction *ϕ*_*r*_ [19]. Therefore, we assume that the fraction of ribosomes translating themselves *χ*_*r*_ is constant for simplicity. To model osmoregulation, we introduce a coupling between the fraction of ribosomes translating the osmolyte-producing protein *χ*_*a*_ and the degree of intracellular crowding. We quantify the crowding effects by the protein density, defined as *ρ*_*p*_ = *m*_*p*_*/V*_*f*_, which serves as a good proxy for the dry-mass density measured in the experiments [43, 44] (see Table 1 and the detailed discussion on the relations between the two densities in Section A of Supplementary Material) and propose the following relation:

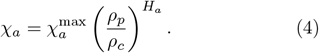

Here, the parameter *H*_*a*_ quantifies the sensitivity of osmoregulation to intracellular crowding. *ρ*_*c*_ is the critical protein density above which intracellular processes are frozen, which we introduce later in Eq. (8). Therefore, 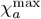 represents the largest possible *ϕ*_*a*_ since all intracellular dynamics is frozen when *ρ*_*p*_ *> ρ*_*c*_. We remark that our model can be directly generalized to cases where osmolyte molecules are directly extracted from the environment. One only needs to change the interpretation of the parameter *k*_*a*_ in Eq. (17) from the synthesis rate to the uptake rate, and all the results are the same.

#### Cell-wall synthesis regulation

In this work, the cell wall is regarded as a linear elastic material, where the turgor pressure is proportional to the elastic strain of the cell wall by a constant modulus *G* such that

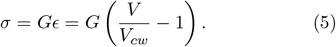

Here, *V*_*cw*_ is the relaxed cell-wall volume (Figure 1C). When plasmolysis happens, the cell membrane detaches from the cell wall (*V < V*_*cw*_), and the turgor pressure is zero. We introduce the growth rate of the relaxed cell-wall volume as 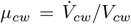.Given that in the steady states of cell growth, *µ*_*r*_ = *µ*_*cw*_, we write *µ*_*cw*_ in the following form without loosing generality,

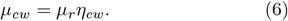

Here, *η*_*cw*_ is a coarse-grained parameter modeling the active regulation of cell-wall synthesis, which we refer to as the cell-wall synthesis efficiency in the following.

Experiments suggested that the turgor pressure induce cell-wall synthesis, e.g., through mechanosensors on cell membrane in *S. pombe* [45, 46], by increasing the pore size of the peptidoglycan network [5], and by accelerating the moving velocity of the cell-wall synthesis machinery in *E. coi* [31]. Guided by these ideas, we model the effects of turgor pressure on the time-dependence of the cell-wall synthesis efficiency as

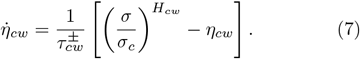

Here, *σ*_*c*_ is a characteristic scale of turgor pressure depending on species. 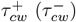 is the relaxation timescale when the current *η*_*cw*_ is below (above) its target value 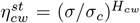.The former (latter) happens immediately after the cell is subject to a hypoosmotic (hyperosmotic) shock. In the extreme case of plasmolysis, the insertion of newly synthesized cell wall materials is interrupted immediately due to the separation of the cell membrane and cell wall. Meanwhile, the up-regulation of cell-wall synthesis rate presumably takes a longer time. For example, in fungi, where polarized growth is generally adopted, the up-regulation of the cell-wall synthesis rate involves reorienting the polarisome complex to the growing tip, directing actin polarization, and delivering cell-wall synthesis machinery [47, 48]. Therefore, we set 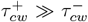 in this work (see details of parameter values in Table 1).

#### Intracellular crowding

Multiple experiments suggested the cytoplasm of bacteria, yeast, and mammalian cells resemble crowded colloidal suspensions in which the mobilities of biomolecules are significantly reduced compared with dilute solutions [14, 15, 49–51], a signature of glass transition [52]. Intracellular crowding affects biochemical processes globally, e.g., slowing down translation and intracellular signaling by suppressing protein diffusion [14, 15, 18, 19, 49]. Therefore, the speed of osmolyte production, translational elongation, and cell-wall synthesis are all slowed down by the same crowding factor:

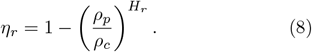

Here, *ρ*_*c*_ is the critical protein density, and *H*_*r*_ is a parameter to quantify the sensitivity of biochemical reactions to the intracellular density. For example, the translational elongation speed is suppressed by intracellular crowding through 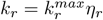.Therefore, the dry-mass growth rate becomes 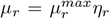,where we introduce 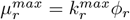.

The details of our model are summarized in Methods, with five independent variables: the protein density *ρ*_*p*_, the mass fraction of osmolyte-producing protein *ϕ*_*a*_, the internal osmotic pressure Π_*in*_, the cell-wall strain *ϵ* and the cell-wall synthesis efficiency *η*_*cw*_. For convenience, Table S2 of the Supplementary Material provides a comprehensive list of all symbols used in the main text along with their meanings.

### Steady states in constant environments

When cell growth reaches a steady state, the proportions of all components, including free water volume, cell mass, and cell wall volume, must be constant relative to the total cell volume to ensure homeostasis. Therefore, all growth rates in steady states of cell growth must be the same: *µ*_*f*_ = *µ*_*r*_ = *µ*_*cw*_. The consequence of cell-wall synthesis regulation can be seen directly from *µ*_*cw*_ = *µ*_*r*_: the turgor pressure at steady states is constant, *σ* = *σ*_*c*_. Experimentally, the cell-wall strain was measured by applying an acute hyperosmotic shock to induce plasmolysis, and it is approximately constant as the external osmolarity increases [22, 53], suggesting a constant turgor pressure independent of external osmolarity, in concert with our model assumptions. The internal osmotic pressure at steady states is related to the external osmotic pressure through Eq. (2),

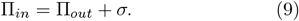

Here, we have neglected the term *µ*_*f*_ */k*_*w*_. Experimentally, an abrupt water flux occurs within hundreds of milliseconds after an osmotic shock [54], from which we can estimate the water permeability as *k*_*w*_ ∼ 100 min^−1^atm^−1^ considering an osmotic shock with an amplitude ΔΠ_*out*_ = 1 atm. Because the typical doubling times of microorganisms are from about 20 minutes to several hours, we estimate *µ*_*f*_ */k*_*w*_ ∼ 10 −100 Pa [54, 55], negligible compared with the typical cytoplasmic osmotic pressures, which can be several atmospheric pressures.

In steady states, the internal osmotic pressure is independent of time. Combining Eq. (4) and the dynamics of the internal osmotic pressure, Eq. (18c), we find the relationships between the protein density, the internal osmotic pressure, and the growth rate in the steady states:

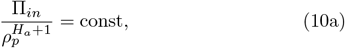

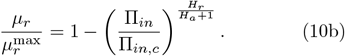

The right-hand side of Eq. (10a) is a constant independent of external osmolarity (see its detailed expression in Section C of Supplementary Material). In deriving Eq. (10b), we have replaced *ρ*_*p*_ by Π_*in*_ in Eq. (8) using Eq. (10a) with the critical internal osmotic pressure Π_*in*,*c*_ proportional to *ρ*_*c*_. Intriguingly, the relationship between the normalized growth rate 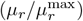 and the normalized cytoplasmic osmotic pressure (Π_*in*_*/*Π_*in*,*c*_), which we refer to as the growth curve in the following, has only one parameter *H*_*r*_*/*(*H*_*a*_ + 1). Therefore, the growth curves of different organisms can be unified by a single formula, Eq. (10b), and different organisms may have different values of *H*_*r*_*/*(*H*_*a*_ + 1). Furthermore, Eq. (10b) predicts a critical external osmolarity Π_*out*,*c*_ = Π_*in*,*c*_ −*σ*_*c*_, beyond which cell growth is completely inhibited.

We test the validity of Eq. (10b) by fitting it to the experimental growth curves (Figure 2A). To do this, we calculate the internal osmotic pressure using Eq. (9) given the values of the external osmotic pressure and the turgor pressure (Table 1). Intriguingly, the growth curves of multiple species can be well fitted by Eq. (10b), from which we infer the parameters *H*_*r*_*/*(*H*_*a*_ + 1) and Π_*in*,*c*_ (Table 1). We find that budding yeast cells exhibit a notable resilience to high external osmolarities: their Π_*in*,*c*_ value is higher than those of Gram-positive bacteria, *B. subtilis*, and Gram-negative bacteria, *E. coli*. Further, budding yeast cells demonstrate a higher value of *H*_*r*_*/*(*H*_*a*_ + 1), indicating a reduced susceptibility to growth rate reduction when exposed to mild increases in the external osmolarity. Meanwhile, the osmoadaptation capability of *E. coli* depends on the growth media, presumably arising from variations in metabolic fluxes and gene expressions [19, 21, 22].

**FIG. 2.**
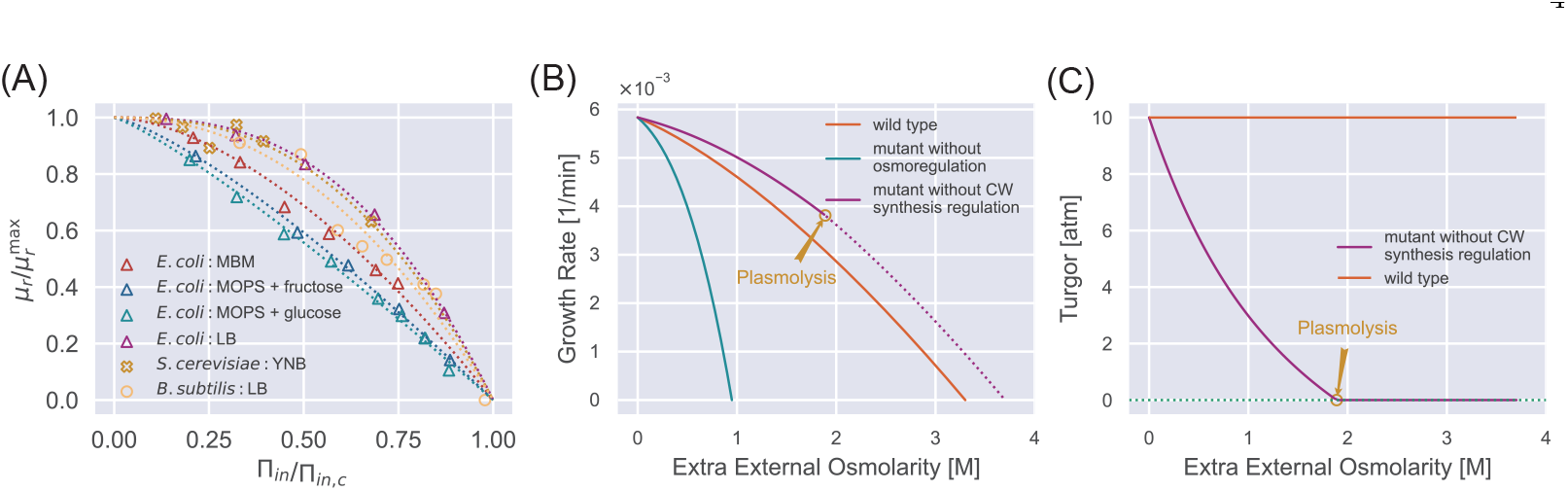
Steady-state properties under a constant external osmolarity. (A) Normalized growth rate vs. normalized internal osmotic pressure of different species under various culture media. The experiment data (scatter markers) are fitted by our theoretical prediction Eq. (10b). The data of *E. coli* are from [19, 21, 22], the data of *B. subtilis* is from [23], and the data of *S. cerevisiae* is from our own experiments. (B) Growth curves of WT cells, mutant cells without osmoregulation (*H*_*a*_ = 0), and mutant cells without cell-wall synthesis regulation (*H*_*cw*_ = 0). The dotted line indicates the region where plasmolysis occurs for the mutant cells with *H*_*cw*_ = 0. (C) Mutant cells without cell-wall synthesis regulation cannot maintain a stable turgor pressure in a hypertonic environment, while WT cells can maintain a constant turgor pressure. The mutant cells reach plasmolysis at a threshold of external osmolarity. In (B) and (C), the parameters for WT cells are chosen as the values for *S. pombe*, and the mutant values are set such that they have the same growth rate as the WT cells in the reference medium (Table S2).

To further reveal the biological functions of biological regulations, we study the steady-state properties of mutant cells in which either osmoregulation or cell-wall synthesis regulation is depleted. For mutant cells without osmoregulation, *H*_*a*_ = 0 in Eq. (4). In this case, the fraction of osmolyte-producing protein is constant with time, i.e., 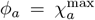.Comparing the dynamics of osmolyte and total protein mass, 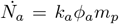 and 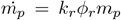,one finds that the ratio of the number of osmolyte molecule and the total protein mass remains constant over time, irrespective of variations in external osmolarity (see the detailed derivation in Section C of Supplementary Material). As the external osmolarity increases, the protein density of mutant cells quickly reaches the critical value *ρ*_*c*_ according to Eq. (10a) with *H*_*a*_ = 0. Therefore, the steady-state growth curve of the mutant cells terminates at an external osmolarity much smaller than wild-type (WT) cells (Figure 2B), in agreement with previous experiments [56].

For mutant cells without the cell-wall synthesis regulation, *H*_*cw*_ = 0; therefore, the cell-wall synthesis efficiency *η*_*cw*_ equals 1 independent of time. Thus, the growth rate of the relaxed cell-wall volume is always equal to the growth rate of total protein mass [Eqs. (6, 7)]. Interestingly, in this case, the turgor pressure at steady states decreases with the increase of external osmolarity (Figure 2C and see the detailed proof in Section C of Supplementary Material). The decreased turgor pressure lowers the internal osmotic pressure given the same Π_*out*_ according to Eq. (9), leading to a lower protein density of mutant cells than WT cells according to Eq. (10a). Therefore, mutant cells grow faster than WT cells under the same external osmolarity (Figure 2B). Nevertheless, the mutant cells are prone to plasmolysis at a threshold external osmolarity where the WT cells can maintain constant turgor pressure (see the vertical line in Figure 2C around 2 M extra external osmolarity). Reduced turgor pressure is detrimental to multiple biological processes, e.g., cytokinesis in fission yeast requires the participation of turgor pressure [57].

To summarize, osmoregulation allows cells to grow in a wide range of external osmolarity conditions with a mild change in protein density. The cell-wall synthesis regulation allows cells to maintain a stable turgor pressure and avoid plasmolysis. Both regulatory mechanisms expand the range of external osmolarities that cells can adapt to.

### Transient dynamics after a constant osmotic shock

Next, we study the dynamical behaviors of cellular properties in response to a constant osmotic shock: the external osmolarity changes abruptly and keeps its value for an infinitely long time. Intriguingly, we find that the dynamics of osmoresponse can be split into shock and adaptation periods (see insets of Figure 3C, D). The immediate water flow due to osmotic imbalance occurs in the shock period, during which the mass and osmolyte productions are negligible. Therefore, the ratio of the internal osmotic pressure and the protein density is invariant right before and after a shock period: 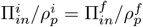 where the upper index *i* (*f*) means the state right before (after) the shock period. Given this condition, we introduce the normalized protein density 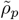 as

**FIG. 3.**
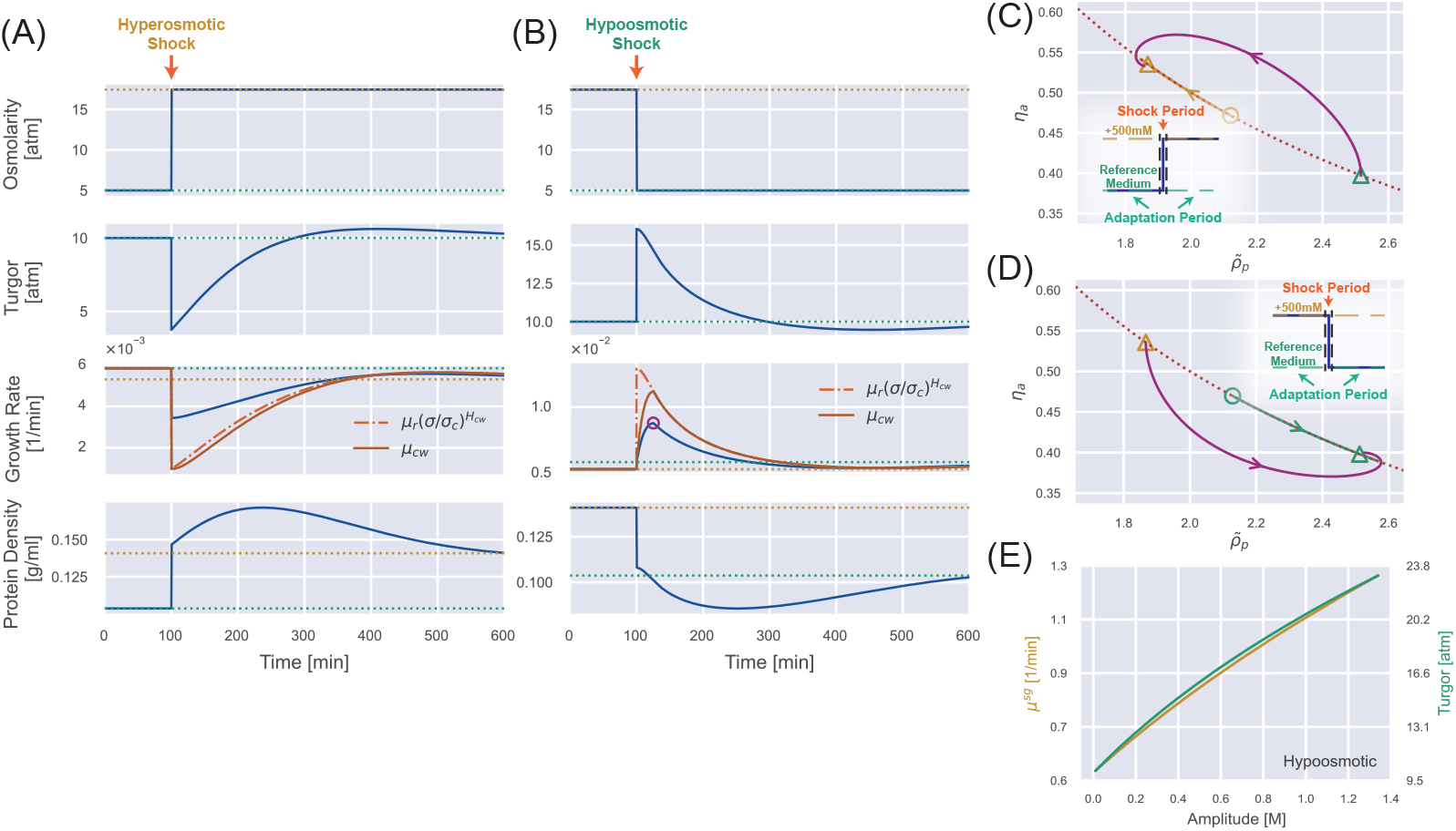
Transient dynamics after a constant osmotic shock. (A) Numerical simulations of cells undergoing a constant 500 mM hyperosmotic shock. The dotted lines represent the steady-state values for the reference growth medium (green) and the medium after perturbation (yellow). (B) Numerical simulations of cells undergoing a constant 500 mM hypoosmotic shock. The purple circle in the third panel marks the growth rate peak during the supergrowth phase. (C) The dynamics of the internal state of a cell characterized by 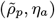.The dotted curve represents the constraint on the steady-state solution 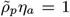,and the solid trajectory is from numerical simulations. The triangles indicate the steady-state solution before the perturbation and the steady-state solution after the perturbation for a long enough time. The yellow open circle represents the immediate steady-state solution after applying the hyperosmotic shock. (D) The same analysis as (C) but for a constant 500 mM hypoosmotic shock. (E) The growth rate peak in the supergrowth phase (yellow) and the immediate value of turgor pressure after the hypoosmotic shock *σ*^*f*^ (green) vs. the amplitude of the hypoosmotic shock.

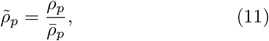

where the normalization factor 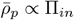 (see its detailed expression in Methods) so that 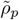 changes continuously across the shock period. Interestingly, we find that osmoresponse are governed by a two-dimensional dynamical system composed of 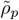 and 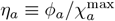 (Methods):

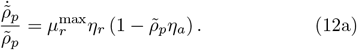

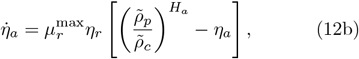

Here, 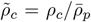 is the normalized critical protein density, and *η*_*a*_ denotes the efficiency of osmoregulation. From the above equations, it is clear that the timescale of osmoregulation is set by the doubling time: it takes about the doubling time for the protein density and the fraction of osmolyte-producing protein to adapt to the new steady-state values. For walled cells, 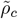 and 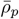 depend on the time since Π_*in*_ = Π_*out*_ + *σ* and the turgor pressure *σ* is time-dependent during osmoresponse processes (Figure 3A, B). For unwalled cells, such as mammalian cells and microbial cells with cell walls removed (i.e., protoplasts), 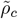 is constant in a fixed environment (see detailed discussion on the transient dynamics of unwalled cells in Section D of Supplementary Material).

Upon a constant hyperosmotic shock, the immediate water efflux leads to an instantaneous drop in turgor pressure and a rise in protein density (Figure 3A). The internal state of the cell, 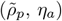 evolves towards to the new equilibrium point, 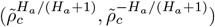 One should note that the equilibrium point is time-dependent initially but eventually becomes fixed as the turgor pressure relaxes to the steady-state value (Figure 3C and Movie S1). Interestingly, the protein density increases initially and then decreases after the shock (Figure 3A). The decrease in protein density is because of the osmoregulation process, which is set by the doubling time [Eqs. (12a, 12b)]. Meanwhile, we find that the initial increase of protein density is because of the suppressed growth of the relaxed cell-wall volume due to the low turgor pressure. Indeed, for unwalled cells, the protein density *ρ*_*p*_ decreases immediately after the shock (Figure S2B). We note that the growth rate approaches the new steady-state value non-monotonically (Figure 3A) because of the spiral trajectory in the space of the internal state (Figure 3C), consistent with experimental observations [22].

The phenomena are essentially the opposite for a constant hypoosmotic shock (Figure 3B, D and Movie S2). However, we find an extremely fast cell growth after the hypoosmotic shock, with a growth rate peak occurring about 25 minutes after applying the shock, which we call the supergrowth phase [27]. One should note that 25 minutes is much shorter than the doubling time (about two hours) but comparable to the timescale of cell-wall synthesis regulation, which we set as 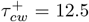 min in the simulations in Figure 3 (we will explain why we choose 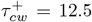 min in the next section). Furthermore, applying a hypoosmotic shock to an unwalled cell does not induce a significant supergrowth phase compared with walled cells (Figure S2D).

We propose that supergrowth comes from the high turgor pressure caused by the hypoosmotic shock, which leads to a fast cell-wall synthesis according to Eq. (7). Rapid insertion of materials into the cell wall relaxes the turgor pressure and allows the cells to grow faster [Eqs. (2, 3)]. This idea is consistent with the observation that the growth rate and the growth rate of the relaxed cell-wall volume *µ*_*cw*_ reach their peaks simultaneously (Figure 3B). This observation also suggests that the timescale of supergrowth, i.e., the timing of growth rate peak, is set by the time scale of cell-wall synthesis regulation [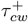 in Eq. (7)]. Notably, in the initial stage of the adaptation period, *µ*_*cw*_ approaches its target from below and reaches its target value at the growth rate peak (i.e., 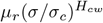) (the third panel of Figure 3B), after which *µ*_*cw*_ sticks to its target value and decreases accordingly because of the short relaxation time 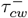 [Eq. (7)]. A detailed proof of the conditions for supergrowth, including the necessity of a cell wall and the regulation of cell-wall synthesis, is provided in Section E of the Supplementary Material.

Following the discussion above, we obtain an analytical expression of the growth rate peak after a hypoosmotic shock (see the detailed derivations in Section F of Supplementary Material)

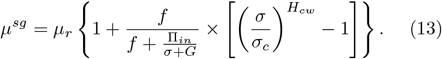

Here, all the variables on the right side are at the growth rate peak. Because the timescale of the osmoresponse process, which is around hours (Figure 3B), is much longer than the timescale of the supergrowth phase, which is about 20 minutes for *S. pombe* [27], the turgor pressure at the growth rate peak can be well approximated by its immediate value after the shock. Therefore, the growth rate peak must increase as the amplitude of the hypoosmotic shock increases, which we confirm numerically (Figure 3E).

### Comparison with experiments: supergrowth phenomena after osmotic oscillation

Next, we quantitatively compare our theoretical predictions regarding the supergrowth phase with experimental data. In Ref. [27], Knapp et al. applied an osmotic oscillation to fission yeast *S. pombe* during which the external osmolarity alternated between two values.

They found cell growth was almost inhibited during the perturbation, while the protein and dry-mass densities increased. Surprisingly, cells grew unusually fast after the osmotic oscillation was removed and reached their maximum growth rate about 20 minutes after the end of the osmotic oscillation. The maximum growth rate can be twice the growth rate in the reference growth medium, and the elevation in growth rate can persist for 2-3 cell cycles. These observations are very similar to our results for a constant hypoosmotic shock (Figure 3B).

To test if our osmoresponse model captures the supergrowth phase for a periodic perturbation, we simulate a wide-type *S. pombe* cells with the same protocols as the experiments (see details of simulations in Methods). Intriguingly, our model successfully recapitulates the supergrowth phase and the gradually increasing protein density and dry-mass density during the perturbation (Figure 4A). We confirm that the cell-wall synthesis regulation is crucial for the emergence of the supergrowth phase since unwalled cells do not exhibit supergrowth under the same conditions (Figure S3B). Interestingly, we find that an infinitely long periodic osmotic shock can be equivalently mapped to a constant osmotic shock (see the detailed discussions and proof in Section D of Supplementary Material), which means that they have the same time-averaged growth rate and protein density in the steady states (Figure S2F).

**FIG. 4.**
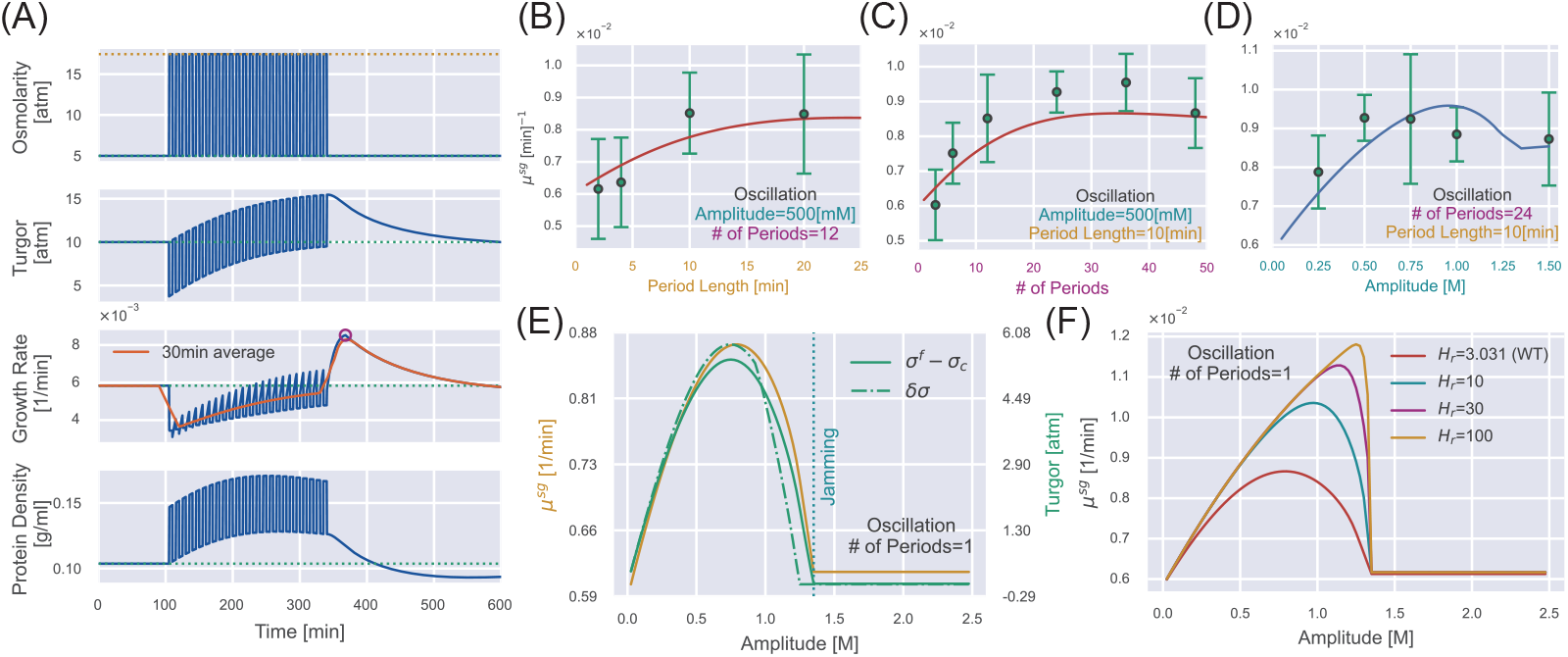
supergrowth phenomena under osmotic oscillation. (A) Numerical simulations of WT *S. pombe* undergoes 24 cycles of 500 mM osmotic oscillations with a 10-minute period. We show a 30-minute window average in the third panel of growth rate. (B-D) Quantitative agreement between simulations and experiments for the growth rate peak *µ*^*sg*^ vs. different oscillation parameters, including (B) amplitude, (C) period length, and (D) number of periods. The red lines in (B, C) are predictions, and the blue line in (D) is fitting from which we infer the values of *H*_*cw*_ and 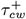.Green dots with error bars are experimental data from Ref. [27]. (E) In the case of osmotic oscillation with a single period, the hyperosmotic period persists for 120 min before reverting to the reference medium. The vertical dotted blue line represents the minimal amplitude to induce cytoplasm jamming during the hyperosmotic period. The excess turgor pressure *σ*^*f*^ −*σ*_*c*_ upon exiting the hyperosmotic period is approximately equal to the recovered turgor pressure *δσ* during the hyperosmotic period. (F) The growth rate peak *µ*^*sg*^ at different *H*_*r*_ vs. the amplitude of a single oscillation. *H*_*r*_ = 3.031 is the value of the WT *S. pombe*. Parameters of WT *S. pombe* are used in this figure unless otherwise mentioned (Table 1).

In Ref. [27], the authors measured the growth rate peaks vs. three different parameters of the osmotic oscillations: amplitude, period length, and number of periods. We first fit the growth rate peaks vs. the amplitudes (Figure 4D), from which we obtain *H*_*cw*_ = 1.7, the sensitivity of the cell-wall synthesis efficiency to turgor pressure [Eq. (7)], and 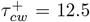 min, the timescale in the up-regulation of cell-wall synthesis efficiency (which is why we set 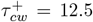 min in the previous section). Other model parameters are inferred from independent steady-state measurements, and we set the timescale in the down-regulation of cell-wall synthesis efficiency as 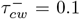 min for simplicity (Table 1). We next fix the values of *H*_*cw*_ and 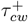 and plot the predicted growth rate peaks vs. the period length (Figure 4B) and number of periods (Figure 4C). As a strong support of our model, our predictions quantitatively match the experimental data without any further fitting.

Two interesting features of the curve *µ*^*sg*^ v.s amplitude catch our attention: the non-monotonic behavior and the kink point at which the derivative is discontinuous (Figure 4D), which are conserved regardless of the number of periods (Figure S7). Therefore, we study the case of a single oscillation for simplicity, which is equivalent to a hyperosmotic shock of finite duration. For a mild hyperosmotic shock, during the period of hyperosmotic shock, the turgor pressure has almost recovered to the steady-state value *σ*_*c*_ (Figure 3A). Therefore, switching from a long hyperosmotic period to the reference growth medium is equivalent to a constant hypoosmotic shock, where we have shown that the growth rate peak increases with the amplitude (Figure 3E). However, the crowding effect becomes more pronounced as the amplitude increases. Beyond the critical amplitude at the kink point, the cytoplasm is completely jammed during the hyperosmotic shock such that the cell states are precisely the same before and after the hyperosmotic shock, which means no supergrowth phase beyond this critical amplitude. Therefore, the curve *µ*^*sg*^ vs. amplitude must be non-monotonic (Figure 4E). Notably, for a very large *H*_*r*_, cells can feel the crowding effect only when the cytoplasm is close enough to the critical protein density, shown as the abrupt decline of *µ*^*sg*^ (Figure 4F).

Finally, we remark that the significance of supergrowth is intimately related to the amount of recovered turgor pressure during the hyperosmotic shock *δσ*. We prove that the overshoot of turgor pressure after the removal of hyperosmotic shock (*σ*^*f*^− *σ*_*c*_), which sets the growth rate peak, is mainly set by the recovered turgor pressure during the hyperosmotic shock (see the detailed discussions in Section G of Supplementary Material). Indeed, *µ*^*sg*^, *σ*^*f*^ −*σ*_*c*_, and *δσ* are highly correlated as we change the amplitude (Figure 4E).

## DISCUSSIONS

This study presents a theory of microbial osmoresponses based on a physical foundation and simplified biological regulation strategies. Our theory captures the steady-state properties of constant turgor pressure and reduced growth rate with increasing external osmolarity. We remark that the growth rate reduction is due to the loss of free water and subsequent intracellular crowding as the external osmolarity increases. In particular, we predict a critical external osmolarity above which cell growth is completely inhibited and a universal relationship between the normalized growth rate and the normalized internal osmotic pressure, fitting the data of bacteria and yeast. We also demonstrate the biological functions of osmoregulation and cell-wall synthesis regulation. Cells defective in osmoregulation cannot grow even if the external osmolarity is only mildly higher than the reference value. Cells defective in cell-wall synthesis regulation cannot maintain turgor pressure as the external osmolarity increases even though they grow faster than WT cells (Figure 2B), which will be a strong support of our theory if confirmed by experiments.

Regarding dynamic behaviors, our model predicts a non-monotonic time dependence of protein density after a constant hyperosmotic shock. We also unveil the supergrowth phase after a hypoosmotic shock, initially discovered in fission yeast after an osmotic oscillation [27]. As a strong support of our theory, the predicted growth rate peaks quantitatively agree with the experimental data without additional fitting. We demonstrate the critical role of cell-wall synthesis regulation in the supergrowth phenomenon (Section E of Supplementary Material). In Ref. [27], the authors observed the rapid repolarization of the cell-wall glucan synthase Bgs4 to the cell tip following the removal of osmotic oscillations in fission yeast, in agreement with the dynamics of the cell-wall synthesis efficiency predicted from our model (compare Figure S11 in this work and Figure S4H in Ref. [27]). To test our theories, we propose applying a hyperosmotic shock with a finite duration and measuring the growth rate after removing the hyperosmotic shock. We predict that the growth rate peak during the supergrowth phase is a nonmonotonic function of shock amplitude, initially rising because of the increased excess turgor pressure and later declining because the protein density reaches the critical value *ρ*_*c*_ during the shock (Figure 4E).

We remark that our model is intrinsically a coarsegrained model with many molecular details regarding gene expression regulation neglected, which allows us to gain more analytical insights. In Ref. [58], the authors studied the responses to osmotic stress in glucose-limited environments and found that cells exhibited stronger osmotic gene expression response under glucose-limited conditions than under glucose-rich conditions. Using a computational model based on molecular mechanisms combined with experimental measurements, the authors demonstrated that in a glucose-limited environment, glycolysis intermediates were limited, which required cells to express more glycerol-production enzymes for stress adaptation. In the current version of our model, we do not account for the interaction between cell growth and osmolyte production; instead, we assume a constant fraction of ribosomes dedicated to translating ribosomal proteins. Our model can be further generalized to include the more complex interactions, including the coupling between biomass and osmolyte production, e.g., by allowing the fraction of ribosomes translating (*χ*_*r*_) to depend on the translation strategy of the osmolyte-producing enzyme (*χ*_*a*_).

Ref. [22] showed that the expansion of *E. coli* cell wall is not directly regulated by turgor pressure, and this scenario is also included in our model as the case of *H*_*cw*_ = 0. According to our model, the supergrowth phase is absent if *H*_*cw*_ = 0 (Figure S8), consistent with the absence of a growth rate peak after a hypoosmotic shock in the experiments of *E. coli* [22]. Meanwhile, our predictions are consistent with the growth rate peak after a hypoosmotic shock observed for *B. subtilis* [23].

We remark several limitations of our current coarsegrained model. First, the high membrane tension that inhibits transmembrane flux of peptidoglycan precursors, leading to a growth inhibition before the supergrowth peak [23] is beyond our model. Second, in our current framework, the osmoregulation and cell-wall synthesis regulation rely on the instantaneous cellular states. However, microorganisms can exhibit memory effects to external stimuli by adapting to their temporal order of appearance [59]. Notably, in the osmoregulation of yeast, a short-term memory, facilitated by post-translational regulation of the trehalose metabolism pathway, and a longterm memory, orchestrated by transcription factors and mRNP granules, have been identified [60]. Besides, our model does not account for the role of osmolyte export in osmoregulation [61] and the interaction between biomass and osmolyte production [58]. Extending our model to include more realistic biological processes will be interesting.

In this work, we construct a systems-level and coarsegrained model capable of elucidating the complex processes underlying microbial osmoresponse. By leveraging the separation of timescales associated with mechanical equilibrium, cell-wall synthesis regulation, and osmoregulation, our model facilitates in-depth analytical and numerical analysis of how these various processes interact during cellular adaptation. In particular, we demonstrate the key physiological functions of osmoregulation and cell-wall synthesis regulation. We then apply this model to interpret the unusual phenomenon of supergrowth observed in fission yeast. This application addresses an essential challenge in experimental studies: exclusive knockout experiments can be difficult, and mechanistic interpretations of experimental observations are often lacking. Our theoretical framework offers a valuable tool for understanding such phenomena, contributing to the fundamental knowledge of microbial physiology and developing predictive models for microbial behavior under osmotic stress.

## ACKNOWLEDGMENTS

We thank Chunxiong Luo for helpful discussions related to this work. The research was funded by the National Key R&D Program of China (2021YFF1200500) and supported by Peking-Tsinghua Center for Life Sciences grants.

## METHODS

### Details of the osmoresponse model

We define the fractions of osmolyte-producing protein and ribosomal proteins in the total proteome as *ϕ*_*a*_ = *m*_*p*,*a*_*/m*_*p*_ and *ϕ*_*r*_ = *m*_*p*,*r*_*/m*_*p*_, respectively. To model gene expression regulation, we introduce *χ*_*a*_ and *χ*_*r*_ as the fractions of ribosomes translating the osmolyte-producing protein and ribosomal proteins such that

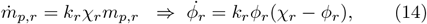

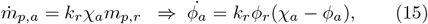

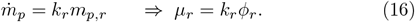

Here, *k*_*r*_ is proportional to the elongation speed of ribosomes on mRNAs divided by the protein mass of a single ribosome, which is affected by the global crowding effect as 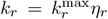.Here, *µ*_*r*_ is the growth rate of total protein mass, which is also the growth rate of dry mass and bound volume in our model since they are all proportional.

The osmolyte molecules are produced by the osmolyte-producing protein, with the rate given by

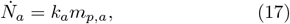

where 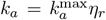 is the osmolyte production rate including the crowding factor and *m*_*p*,*a*_ is the mass of osmolyte-producing protein. We summarize the dynamical equations involved in the osmoresponse model:

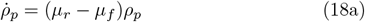

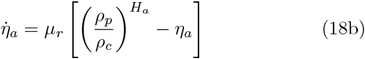

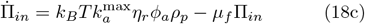

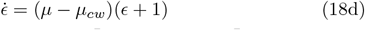

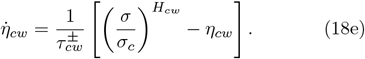

To describe the osmoregulation process using a two-dimensional dynamical system, we introduce the normalized protein density as

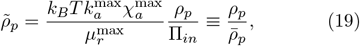

Combining Eq. (11) and Eq. (18a), we obtain the dynamical equation for 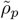 as

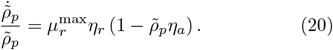

Using Eq. (15), we obtain the equation for 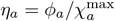 as

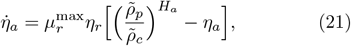

where 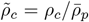.The unique equilibrium point for the internal state is

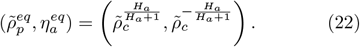

### Details of numerical simulations

We employ the LSODA algorithm with automatic stiffness detection and switching [63], implemented in SciPy [64], to solve Eq. (18). The parameters used for numerical simulations of walled cells are listed in Table 1.

## References

[1] L. N. Csonka, Physiological and genetic responses of bacteria to osmotic stress, Microbiological Reviews 53, 121 (1989).

[2] D. Muzzey, C. Gómez-Uribe, J. Mettetal, and A. Oudenaarden, A systems-level analysis of perfect adaptation in yeast osmoregulation, Cell 138, 160 (2009).

[3] J. M. Wood, Bacterial responses to osmotic challenges, Journal of General Physiology 145, 381 (2015).

[4] E. Bremer and R. Krämer, Responses of microorganisms to osmotic stress, Annual Review of Microbiology 73, 313 (2019), pMID: 31180805.

[5] A. Typas, M. Banzhaf, B. van den Berg van Saparoea, J. Verheul, J. Biboy, R. J. Nichols, M. Zietek, K. Beilharz, K. Kannenberg, M. von Rechenberg, E. Breukink, T. den Blaauwen, C. A. Gross, and W. Vollmer, Regulation of peptidoglycan synthesis by outer-membrane proteins, Cell 143, 1097 (2010).

[6] E. R. Rojas and K. C. Huang, Regulation of microbial growth by turgor pressure, Current Opinion in Microbiology 42, 62 (2018), cell Regulation.

[7] C. Cadart, L. Venkova, P. Recho, M. C. Lagomarsino, and M. Piel, The physics of cell-size regulation across timescales, Nature Physics 15, 993 (2019).

[8] B. Kempf and E. Bremer, Uptake and synthesis of compatible solutes as microbial stress responses to high-osmolality environments, Archives of Microbiology 170, 319 (1998).

[9] M. Burg, Macromolecular crowding as a cell volume sensor, Cellular physiology and biochemistry : international journal of experimental cellular physiology, biochemistry, and pharmacology 10, 251 (2000).

[10] J. Berg, A. Boersma, and B. Poolman, Microorganisms maintain crowding homeostasis, Nature Reviews Microbiology 15 (2017).

[11] M. Model, J. Hollembeak, and M. Kurokawa, Macromolecular crowding: a hidden link between cell volume and everything else, Cellular physiology and biochemistry : international journal of experimental cellular physiology, biochemistry, and pharmacology 55, 25 (2021).

[12] J. A. Dix and A. Verkman, Crowding effects on diffusion in solutions and cells, Annual Review of Biophysics 37, 247 (2008), pMID: 18573081.

[13] K. A. Dill, K. Ghosh, and J. D. Schmit, Physical limits of cells and proteomes, Proceedings of the National Academy of Sciences 108, 17876 (2011).

[14] A. Miermont, F. Waharte, S. Hu, M. N. McClean, S. Bottani, S. Léon, and P. Hersen, Severe osmotic compression triggers a slowdown of intracellular signaling, which can be explained by molecular crowding, Proceedings of the National Academy of Sciences 110, 5725 (2013).

[15] J. T. Mika, P. E. Schavemaker, V. Krasnikov, and B. Poolman, Impact of osmotic stress on protein diffusion in lactococcus lactis, Molecular Microbiology 94, 857 (2014).

[16] M. C. Munder, D. Midtvedt, T. Franzmann, E. Nuske, O. Otto, M. Herbig, E. Ulbricht, P. Müller, A. Taubenberger, S. Maharana, et al., A ph-driven transition of the cytoplasm from a fluid-to a solid-like state promotes entry into dormancy, eLife 5, e09347 (2016).

[17] R. P. Joyner, J. H. Tang, J. Helenius, E. Dultz, C. Brune, L. J. Holt, S. Huet, D. J. Müller, and K. Weis, A glucose-starvation response regulates the diffusion of macro-molecules, eLife 5, e09376 (2016).

[18] A. T. Molines, J. Lemière, M. Gazzola, I. E. Steinmark, C. H. Edrington, C.-T. Hsu, P. Real-Calderon, K. Suhling, G. Goshima, L. J. Holt, et al., Physical properties of the cytoplasm modulate the rates of microtubule polymerization and depolymerization, Developmental Cell 57, 466 (2022).

[19] X. Dai, M. Zhu, M. Warren, R. Balakrishnan, H. Okano, J. R. Williamson, K. Fredrick, and T. Hwa, Slowdown of translational elongation in escherichia coli under hyperosmotic stress, mBio 9, e02375 (2018).

[20] Y. Chen, J.-H. Huang, C. Phong, and J. James E. Ferrell, Protein homeostasis from diffusion-dependent control of protein synthesis and degradation, bioRxiv 10.1101/2023.04.24.538146 (2023).

[21] S. Cayley, B. A. Lewis, H. J. Guttman, and M. T. Record Jr, Characterization of the cytoplasm of escherichia coli k-12 as a function of external osmolarity: implications for protein-dna interactions in vivo, Journal of molecular biology 222, 281 (1991).

[22] E. Rojas, J. A. Theriot, and K. C. Huang, Response of escherichia coli growth rate to osmotic shock, Proceedings of the National Academy of Sciences 111, 7807 (2014).

[23] E. R. Rojas, K. C. Huang, and J. A. Theriot, Homeostatic cell growth is accomplished mechanically through membrane tension inhibition of cell-wall synthesis, Cell systems 5, 578 (2017).

[24] W. Scott, Water relations of staphylococcus aureus at 30 c, Australian journal of biological sciences 6, 549 (1953).

[25] J. Christian and W. Scott, Water relations of salmonellae at 30 c, Australian journal of biological sciences 6, 565 (1953).

[26] J. Christian, The influence of nutrition on the water relations of salmonella oranienburg, Australian Journal of Biological Sciences 8, 75 (1955).

[27] B. D. Knapp, P. Odermatt, E. R. Rojas, W. Cheng, X. He, K. C. Huang, and F. Chang, Decoupling of rates of protein synthesis from cell expansion leads to supergrowth, Cell Systems 9, 434 (2019).

[28] R. Rollin, J.-F. Joanny, and P. Sens, Physical basis of the cell size scaling laws, Elife 12, e82490 (2023).

[29] M. Scott, C. W. Gunderson, E. M. Mateescu, Z. Zhang, and T. Hwa, Interdependence of Cell Growth and Gene Expression: Origins and Consequences, Science 330, 1099 (2010).

[30] H. Jiang and S. X. Sun, Morphology, growth, and size limit of bacterial cells, Phys. Rev. Lett. 105, 028101 (2010).

[31] A. Amir and D. R. Nelson, Dislocation-mediated growth of bacterial cell walls, Proceedings of the National Academy of Sciences 109, 9833 (2012).

[32] D. S. Cayley, H. J. Guttman, and M. T. Record, Biophysical characterization of changes in amounts and activity of escherichia coli cell and compartment water and turgor pressure in response to osmotic stress, Biophysical journal 78, 1748 (2000).

[33] A. M. Whatmore and R. H. Reed, Determination of turgor pressure in bacillus subtilis: a possible role for k+ in turgor regulation, Microbiology 136, 2521 (1990).

[34] S. Cayley and M. Record, Roles of cytoplasmic osmolytes, water, and crowding in the response of escherichia coli to osmotic stress: Biophysical basis of osmoprotection by glycine betaine, Biochemistry 42, 12596 (2003).

[35] J. Lemière and F. Chang, Quantifying turgor pressure in budding and fission yeasts based upon osmotic properties, bioRxiv 10.1101/2023.06.07.544129 (2023).

[36] E. Zhou, X. Trepat, C. Park, G. Lenormand, M. Oliver, S. Mijailovich, C. Hardin, D. Weitz, J. Butler, and J. Fredberg, Universal behavior of the osmotically compressed cell and its analogy to the colloidal glass transition, Proceedings of the National Academy of Sciences 106, 10632 (2009).

[37] E. I. Solenov, G. S. Baturina, L. E. Katkova, and S. G. Zarogiannis, Methods to measure water permeability., Advances in experimental medicine and biology 969, 263 (2017).

[38] N. Empadinhas and M. da Costa, Osmoadaptation mechanisms in prokaryotes: Distribution of compatible solutes, International microbiology : the official journal of the Spanish Society for Microbiology 11, 151 (2008).

[39] R. H. Reed, J. A. Chudek, R. Foster, and G. M. Gadd, Osmotic significance of glycerol accumulation in exponentially growing yeasts, Applied and Environmental Microbiology 53, 2119 (1987).

[40] S. Hohmann, M. Krantz, and B. Nordlander, Yeast osmoregulation., Methods in enzymology 428, 29 (2007).

[41] A. Blomberg, Yeast osmoregulation – glycerol still in pole position, FEMS Yeast Research 22, 10.1093/femsyr/foac035 (2022), foac035.

[42] Q. Wang and J. Lin, Environment-specificity and universality of the microbial growth law, Communications Biology 5, 891 (2022).

[43] F. Delgado, N. Cermak, V. Hecht, S. Son, Y. Li, S. Knudsen, S. Olcum, J. Higgins, W. Grover, and S. Manalis, Intracellular water exchange for measuring the dry mass, water mass and changes in chemical composition of living cells, PloS one 8, e67590 (2013).

[44] P. D. Odermatt, T. P. Miettinen, J. Lemière, J. H. Kang, E. Bostan, S. R. Manalis, K. C. Huang, and F. Chang, Variations of intracellular density during the cell cycle arise from tip-growth regulation in fission yeast, Elife 10, e64901 (2021).

[45] V. Dupres, D. Alsteens, S. Wilk, B. Hansen, J. J. Heinisch, and Y. F. Dufrêne, The yeast wsc1 cell surface sensor behaves like a nanospring in vivo, Nature chemical biology 5, 857 (2009).

[46] R. Neeli-Venkata, C. M. Diaz, R. Celador, Y. Sanchez, and N. Minc, Detection of surface forces by the cell-wall mechanosensor wsc1 in yeast, Developmental cell 56, 2856 (2021).

[47] K. Kono, Y. Saeki, S. Yoshida, K. Tanaka, and D. Pellman, Proteasomal degradation resolves competition between cell polarization and cellular wound healing, Cell 150, 151 (2012).

[48] A. Haupt, D. Ershov, and N. Minc, A positive feedback between growth and polarity provides directional persistency and flexibility to the process of tip growth, Current Biology 28, 3342 (2018).

[49] B. Parry, I. Surovtsev, M. Cabeen, C. O’Hern, E. Dufresne, and C. Jacobs-Wagner, The bacterial cytoplasm has glass-like properties and is fluidized by metabolic activity, Cell 156, 183 (2014).

[50] K. Nishizawa, K. Fujiwara, M. Ikenaga, N. Nakajo, M. Yanagisawa, and D. Mizuno, Universal glass-forming behavior of in vitro and living cytoplasm, Scientific Reports 7 (2017).

[51] H. Ebata, K. Umeda, K. Nishizawa, W. Nagao, S. Inokuchi, Y. Sugino, T. Miyamoto, and D. Mizuno, Activity-dependent glassy cell mechanics i: Mechanical properties measured with active microrheology, Biophysical Journal 122, 1781 (2023).

[52] G. L. Hunter and E. R. Weeks, The physics of the colloidal glass transition, Reports on Progress in Physics 75, 066501 (2012).

[53] G. Misra, E. R. Rojas, A. Gopinathan, and K. Huang, Mechanical consequences of cell-wall turnover in the elongation of a gram-positive bacterium, Biophysical Journal 104, 2342 (2013).

[54] M. Boer, A. Anishkin, and S. Sukharev, Adaptive mscs gating in the osmotic permeability response in e. coli: the question of time, Biochemistry 50, 4087 (2011).

[55] R. Ye and A. Verkman, Simultaneous optical measurement of osmotic and diffusional water permeability in cells and liposomes, Biochemistry 28, 824 (1989).

[56] J. L. Brewster, T. de Valoir, N. D. Dwyer, E. Winter, and M. C. Gustin, An osmosensing signal transduction pathway in yeast, Science 259, 1760 (1993).

[57] S. A. Proctor, N. Minc, A. Boudaoud, and F. Chang, Contributions of turgor pressure, the contractile ring, and septum assembly to forces in cytokinesis in fission yeast, Current Biology 22, 1601 (2012).

[58] W. Shen, Z. Gao, K. Chen, A. Zhao, Q. Ouyang, and C. Luo, The regulatory mechanism of the yeast osmoresponse under different glucose concentrations, iScience 26, 105809 (2023).

[59] A. Mitchell, G. H. Romano, B. Groisman, A. Yona, E. Dekel, M. Kupiec, O. Dahan, and Y. Pilpel, Adaptive prediction of environmental changes by microorganisms, Nature 460, 220 (2009).

[60] Y. Jiang, Z. AkhavanAghdam, Y. Li, B. M. Zid, and N. Hao, A protein kinase a–regulated network encodes short-and long-lived cellular memories, Science signaling 13, eaay3585 (2020).

[61] M. J. Tamás, K. Luyten, F. C. W. Sutherland, A. Hernandez, J. Albertyn, H. Valadi, H. Li, B. A. Prior, S. G. Kilian, J. Ramos, et al., Fps1p controls the accumulation and release of the compatible solute glycerol in yeast osmoregulation, Molecular microbiology 31, 1087 (1999).

[62] E. Atilgan, V. Magidson, A. Khodjakov, and F. Chang, Morphogenesis of the fission yeast cell through cell wall expansion, Current Biology 25, 2150 (2015).

[63] L. Petzold, Automatic selection of methods for solving stiff and nonstiff systems of ordinary differential equations, SIAM journal on scientific and statistical computing 4, 136 (1983).

[64] P. Virtanen, R. Gommers, T. E. Oliphant, M. Haberland, T. Reddy, D. Cournapeau, E. Burovski, P. Peterson, W. Weckesser, J. Bright, et al., Scipy 1.0: fundamental algorithms for scientific computing in python, Nature methods 17, 261 (2020).

