## Supplementary Material for "Physical constraints and biological regulations underlie universal osmoresponses"

Yiyang Ye,[1] Qirun Wang,[1] and Jie Lin[1,2]
[1] Center for Quantitative Biology, Peking University, Beijing, China
[2] Peking-Tsinghua Center for Life Sciences, Peking University, Beijing, China
(Dated: January 13, 2025)

### A. Conversion between the dry mass density and the protein density

In the main text, we mainly focus on the density of cytoplasmic proteins, defined as $\rho_p = m_p/V_f$, which is related to the dry mass density, defined as $\hat{\rho}_d = m_{tot,d}/V_{tot}$. Here, the subscript $tot$ means the total mass and cell volume, encompassing regions outside the cytoplasm. Taking yeast as an example, the dry mass in the cell wall contributes to approximately 30% of the total dry mass [1], with mannoproteins constituting 40% of the dry mass in the cell wall [2]. Therefore, a nonnegligible proportion of proteins resides outside the cytoplasm in microorganisms. For simplicity, we assume that the fraction of protein mass $m_{tot,p}$ in the total dry mass $m_{tot,d}$ remains constant under different external osmolarities for a given species. Also, this constancy holds both inside and outside the cytoplasm such that

$$\frac{m_p}{m_d} = \frac{m_{tot,p}}{m_{tot,d}} = \text{const.} \tag{S1}$$

Further, we assume that the mass fraction of the cytoplasmic proteins in total proteins is constant, from which we infer the ratio between the total bound volume and the cytoplasmic bound volume: $(V_{tot} - V_f)/V_b = m_{tot,p}/m_p$. With these assumptions, $\rho_p$ and $\hat{\rho}_d$ are related as

$$\rho_p = \frac{m_p}{m_d}\left(\frac{m_p}{m_{tot,p}} + f^{-1} - 1\right)\hat{\rho}_d. \tag{S2}$$

Here, $f$ is the fraction of free volume in the cytoplasmic volume, $f = V_f/V$. In Table S1, we present the dry mass fraction of protein $m_p/m_d$ and the cytoplasmic protein mass fraction $m_p/m_{tot,p}$ used in this work. The same parameters are used for *S. pombe* and *S.cerevisiae* for simplicity. Given Eq. (S2) and the value of $f$, which we discuss in the next section, we obtain the protein density from the dry mass density.

| ***E. coli*** | **Value** | **Reference** |
|---|---|---|
| $m_p/m_{tot,p}$ | 0.8 | Ref. [3] |
| $m_p/m_d$ | 0.68 | Ref. [3] |
| ***S.cerevisiae* & *S. pombe*** | **Value** | **Reference** |
| $m_p/m_{tot,p}$ | 0.65 | Ref. [4] |
| $m_p/m_d$ | 0.4 | Ref. [5] |

TABLE S1. The fraction of protein mass in the total dry mass, $\frac{m_p}{m_d}(=\frac{m_{tot,p}}{m_{tot,d}})$, and the fraction of cytoplasmic protein mass in the total protein mass, $m_p/m_{tot,p}$.

### B. Estimations of the model parameters

In this section, all variables with the superscript $*$ are for the reference growth media we show in Table 1 of the main text.

#### *1. The free volume fraction of S. cerevisiae*

Upon an extreme hyperosmotic shock, the cytoplasm expels all free water such that the expelled volume fraction is equal to the free volume fraction $f$ in the reference growth medium, which is how we compute the free volume fraction for *S. cerevisiae* according to Ref. [6] (Table 1 in the main text).

#### 2. The free volume fraction of *S. pombe*

In Ref. [7], the authors measured the immediate changes in cytoplasmic volume after a constant hyperosmotic shock with varying amplitudes, generating a curve of $V^f/V^*$ vs. $\Delta\Pi_{out}$ where $\Delta\Pi_{out}$ is the extra external osmolarity relative to the reference growth medium. As mentioned in the main text, there is a separation of timescales between the shock periods and the adaptation periods. The number of osmolyte molecules $N_a$ remains conserved during the shock periods. Consequently, the cytoplasmic volume right after the hyperosmotic shock is given by:

$$V^f = V_f^f + V_b^* = \left[\frac{\Pi_{in}^*}{\Pi_{in}^f} f^* + (1 - f^*)\right] V^* \tag{S3}$$

This relationship has been verified for various cell types [8, 9]. Moreover, the relaxed cell-wall volume is also conserved during the shock periods. Using Eq. (S3), the turgor pressure after the shock $\sigma^f$ can be related to the turgor pressure of the reference growth medium by

$$\frac{\sigma^f}{\sigma^*} = 1 + \left(1 + \epsilon^{*-1}\right)\left(\frac{\Pi_{out}^* + \sigma^*}{\Pi_{out}^f + \sigma^f} - 1\right) f^*. \tag{S4}$$

We note that the left-hand side of Eq. (S4) increases monotonically with $\sigma^f$, while the right-hand side decreases monotonically. Therefore, given the values of $\epsilon^*$, $f^*$, and $\sigma^*$, $\sigma^f$ can be solved uniquely from Eq. (S4). In the case of an extreme hyperosmotic shock (i.e., a very large $\Pi_{out}^f$), no positive solution of $\sigma^f$ can be found, corresponding to plasmolysis. After substituting $\Pi_{in}^f = \Pi_{out}^f + \sigma^f$ into Eq. (S3), $V^f/V^*$ is obtained accordingly. Therefore, we infer the values of $f^*$ for *S. pombe* growing in YE5S media using the value of steady-state turgor pressure $\sigma^*$ (Table 1 in the main text) [10], the value of steady-state cell-wall strain $\epsilon^*$ [11], and the data of $V^f/V^*$ vs. $\Delta\Pi_{out}$ (Figure S1).

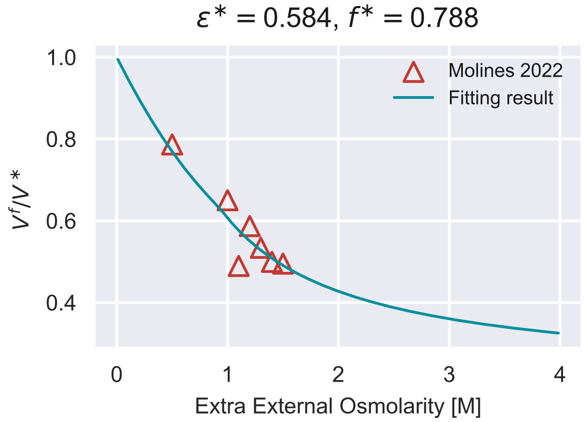


FIG. S1. Fit of $V^f/V^*$ vs. $\Delta\Pi_{out}$ for *S. pombe* growing in YE5S media, from which we infer the value of $f^*$. The data is from Ref. [7]

.

#### 3. Determination of $\rho_c$

In Ref. [7], the authors studied the changes in intracellular biochemical processes and cytoplasmic diffusion coefficients in response to hyperosmotic shock. The authors found that in acute response to hyperosmotic shocks, the interphase microtubule cytoskeleton appeared to freeze at high external osmolarities. Similar phenomena were found when applying a hyperosmotic shock on *S.cerevisiae* [6], where several intracellular signaling cascades, including the stress-response pathways, are significantly slowed down. In our model, the critical external osmolarity $\Pi_{out,c}$ that freezes the cytoplasm happens when $\rho_p = \rho_c$. Therefore, $\rho_c$ can be derived as follows (Table 1 in the main text),

$$\frac{\rho_c}{\rho_p^*} = \frac{\Pi_{out}^* + \sigma^*}{\Pi_{out,c}}. \tag{S5}$$

In the above equation, we have used the fact that the number of osmolyte molecules is conserved during the shock periods and the condition that plasmolysis occurs at the critical external osmolarity such that the turgor pressure is zero under the relevant parameters of *S. cerevisiae* and *S. pombe* (Table 1 in the main text).

#### *4. Determination of* $\alpha$

We define $\alpha$ as the proportionality coefficient between the bound volume $V_b$ and the cytoplasmic protein mass $m_p$, $\alpha = V_b/m_p$. Thus, one can find $\alpha$ as a function of $f^*$ and $\rho_p^*$ through the following equation:

$$\alpha \rho_p^* = \frac{1}{f^*} - 1, \tag{S6}$$

which is how we compute $\alpha$ for *S. cerevisiae* and *S. pombe* (Table 1 in the main text). We remark that the above equation is also valid for non-steady states.

We compute $\alpha$ for *E. coli* alternatively. In Ref. [12], a severe osmotic shock was applied to *E. coli*, squeezing out all of the free water $V_f$, and the remaining volume of bound water per dry mass $V_{bw}/m_{tot,d}$ was computed. Consequently, $\alpha$ can be deduced from the formula:

$$\alpha = \frac{V_{bd}}{m_p} + \frac{V_{bw}}{m_p} = \left( \rho_0^{-1} + \frac{V_{bw}}{m_{tot,d}} \frac{m_{tot,d}}{m_d} \right) \frac{m_d}{m_p}. \tag{S7}$$

Here, $\rho_0 = 1.59$ [g/ml] represents the density of the dry mass, a constant independent of the external osmolarity [3].

### C. Steady-state properties of mutant cells

#### *1. Mutant cells without osmoregulation*

In steady states, the osmoregulation efficiency is fully determined by the cytoplasmic protein density, $\eta_a = \phi_a/\chi_a^{\max} = (\rho_p/\rho_c)^{H_a}$. After substituting $\dot{\Pi}_{in} = 0$ and $\mu_r = \mu_f$ into Eq. (18c) of the main text, we reach the constant shown in Eq. (15a):

$$\frac{\Pi_{in}}{\rho_p^{H_a+1}} = \frac{k_B T k_a^{\max} \chi_a^{\max}}{\rho_c^{H_a} \mu_r^{\max}}. \tag{S8}$$

In the case of the osmoregulation-defective cells, $H_a = 0$, and the osmoregulation efficiency remains constant ($\eta_a = 1$), irrespective of variation in the cytoplasmic protein density $\rho_p$. Considering a steady state denoted by $i$, according to Eq. (S8), the ratio between the internal osmotic pressure and the protein density is

$$\frac{\Pi_{in}^i}{\rho_p^i} = \frac{k_B T k_a^{\max} \chi_a^{\max}}{k_r^{\max} \chi_r} \tag{S9}$$

Combining Eqs. (18a, 18c) and Eq. (S9), we find that the ratio is time-independent even in transient states because

$$\frac{\mathrm{d}}{\mathrm{d}t} \left( \frac{\Pi_{in}}{\rho_p} \right) = \mu_r \left( \frac{\Pi_{in}^i}{\rho_p^i} - \frac{\Pi_{in}}{\rho_p} \right) = 0. \tag{S10}$$

The second equality arises from the argument that at the initial state when $t = 0$, $\Pi_{in}/\rho_p = \Pi_{in}^i/\rho_p^i$. Therefore, $\Pi_{in}/\rho_p = \Pi_{in}^i/\rho_p^i$ establishes during the entire osmoresponses process. Intuitively, this constancy is due to the alignment between the increasing rates of the number of osmolyte molecules and the total protein mass,

$$\dot{N}_a = k_a^{\max} \chi_a^{\max} \eta_r m_p, \tag{S11a}$$
$$\dot{m}_p = k_r^{\max} \chi_r \eta_r m_p. \tag{S11b}$$

The ratio of the two increasing rates is always constant and independent of time.

#### 2. Mutant cells without cell-wall synthesis regulation

For mutant cells with cell-wall synthesis regulation knocked out, the parameter $\mu_{cw}$ is no longer subject to regulation by turgor pressure. This condition is equivalent to setting $H_{cw} = 0$ and $\eta_{cw} = 1$ in our model. Consequently, the cell wall and dry mass productions proceed at the same pace, $\mu_{cw} = \mu_r$. Combing Eq. (18a) and (18d) in the main text, we find the following conserved quantity:

$$\frac{\mathrm{d}}{\mathrm{d}t} \ln \left[ \frac{\epsilon + 1}{\alpha + \rho_p^{-1}} \right] = \frac{\dot{\epsilon}}{\epsilon + 1} + \frac{f \dot{\rho}_p}{\rho_p} = 0. \tag{S12}$$

The relationship between $\alpha$ and $f$ [Eq. (S6)] is utilized in the above derivation. Imagine the external osmotic pressure increases quasistatically, the protein density increases accordingly [Eq. (S8)], leading to a deceased turgor pressure according to Eq. (S12) (Figure 2C in the main text).

The emergence of this conserved quantity can also be seen from the definition of the elastic strain $1 + \epsilon = V/V_{cw}$ and $m_p/V = m_p/(V_b + V_f) = 1/(\alpha + \rho_p^{-1})$, which gives rise to the following relationship:

$$\frac{m_p}{V_{cw}} = \frac{\epsilon + 1}{\alpha + \rho_p^{-1}}. \tag{S13}$$

Since $m_p/V_{cw}$ is constant because $\mu_r = \mu_{cw}$, we obtain Eq. (S12).

### D. Dynamics of unwalled cells

#### 1. Constant osmotic shock

The dynamics of unwalled cells (as described in Eq. (18) of the main text with $\sigma = 0$) consists of three independent variables: $\Pi_{in}$, $\rho_p$ and $\eta_a$. Without turgor pressure, $\Pi_{in} = \Pi_{out}$ holds throughout the adaptation periods. Therefore, the osmoregulation process of unwalled cells can be completely represented by the 2D-trajectory of internal state $(\tilde{\rho}_p, \eta_a)$, where $\tilde{\rho}_p$ is the normalized protein density. Since the normalization factor introduced in Eq. (19) of the main text is solely dependent on the external osmolarity

$$\bar{\rho}_p = \frac{\mu_r^{\max}}{k_B T k_a^{\max} \chi_a^{\max}} \Pi_{out}, \tag{S14}$$

the steady state defined by $\tilde{\rho}_c$ is time-independent for unwalled cell.

In Figure S2A and S2C, we present the osmoresponse processes of unwalled cells to hyper/hypoosmotic shocks in the two-dimensional space of the internal state. Due to the spiral nature of these trajectories, the growth rates converge to new steady-state values in a non-monotonic manner (Figure S2B, D), where the characteristic timescale of the adaptation process is set by the doubling time. Comparing with the response of unwalled cells (Figure 3A, B of the main text), we find that the presence of cell wall buffers the drastic changes in protein density induced by osmotic shocks. In particular, after a constant hypoosmotic shock, the growth rate of unwalled cells decreases immediately without supergrowth, unlike the walled cells shown in Figure 3B of the main text.

#### 2. Oscillatory osmotic perturbation

In Figure S2E, we study the osmoregulation dynamics during an osmotic oscillation. The internal state trajectory approaches different targets during the hyperosmotic and hypoosmotic periods. Eventually, it reaches a periodic steady state around a dynamical equilibrium point on the curve $\tilde{\rho}_p \eta_a = 1$ (Movie S3). In this case, the net changes of $\tilde{\rho}_p$ and $\eta_a$ are zero during one oscillation cycle:

$$\int_{\text{hyper}} \eta_r^+ (1 - \tilde{\rho}_p \eta_a) \, \mathrm{d}t + \int_{\text{hypo}} \eta_r^- (1 - \tilde{\rho}_p \eta_a) \, \mathrm{d}t = 0, \tag{S15a}$$

$$\int_{\text{hyper}} \eta_r^+ \left[ \left( \frac{\tilde{\rho}_p}{\tilde{\rho}_c^+} \right)^{H_a} - \eta_a \right] \mathrm{d}t + \int_{\text{hypo}} \eta_r^- \left[ \left( \frac{\tilde{\rho}_p}{\tilde{\rho}_c^-} \right)^{H_a} - \eta_a \right] \mathrm{d}t = 0. \tag{S15b}$$

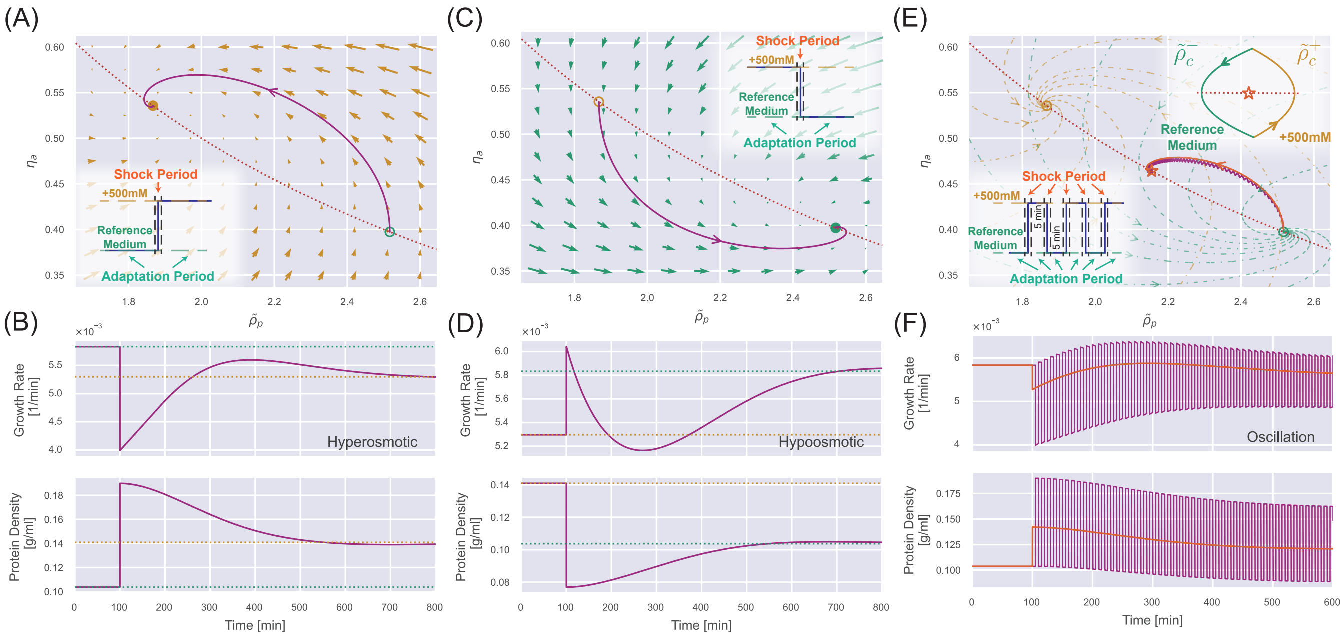


FIG. S2. **Dynamics of osmoresponse for unwalled cells.** (A) The dynamics in the internal state space after a constant 500 mM hyperosmotic shock. The dotted curve represents $\tilde{\rho}_p \eta_a = 1$, and the solid trajectory is from numerical simulations. The arrows represent the stream flow described by Eqs. (20, 21) of the main text. The inset depicts the separation of timescale into shock periods and adaptation periods. (B) The temporal trajectory of growth rate (upper panel) and protein density (lower panel) corresponding to the 500 mM hyperosmotic shock in (A). The dashed lines represent the steady-state values in the reference medium (green) and the medium after perturbation (yellow). (C) The same analysis as (A) but for a constant 500 mM hypoosmotic shock. (D) The same analysis as (B) but for a constant 500 mM hypoosmotic shock. (E) The trajectory of the internal state during a 500 mM osmotic oscillation with a 10-minute period (the purple line). The low inset depicts the separation of timescale into shock periods and adaptation periods. The upper inset shows that the trajectory finally converges to a periodic circle around a dynamic equilibrium point (marked by an orange star) on the red dotted curve $\tilde{\rho}_p \eta_a = 1$. The trajectory for a constant hyperosmotic shock with an amplitude of 222 mM (orange curve) almost coincides with the trajectory for the oscillator shock. (F) The purple curves represent the temporal trajectory of growth rate (upper panel) and protein density (lower panel) corresponding to the 500 mM osmotic oscillation in (E). The orange curves represent the corresponding ones for the constant hyperosmotic shock with an amplitude of 222 mM.

Here, the subscripts $\pm$ of $\eta_r$ stand for the hyperosmotic (500 mM) and hypoosmotic (the reference medium) period, respectively. As the external osmolarity switches, a sharp change occurs in $\eta_r = 1 - (\rho_p/\rho_c)^{H_r} = 1 - (\tilde{\rho}_p/\tilde{\rho}_c)^{H_r}$ due to the different $\tilde{\rho}_c$ values in the hyper/hypoosmotic periods.

We consider the limiting case where the oscillation period $T$ approaches 0. In this case, the values of $\tilde{\rho}_p$ and $\eta_a$ along the cycles can be well approximated by their values at the dynamical equilibrium point. Thus, we must have $\tilde{\rho}_p \eta_a = 1$ for the dynamical equilibrium point according to Eq. (S15a). This result suggests that the dynamic equilibrium point also lies on the curve for the steady states.

For unwalled cells, $\tilde{\rho}_c$ maintains a fixed value during the hyper/hypoosmotic period. After applying the same procedure to Eq. (S15b), the following relationship is derived,

$$(\eta_r^+ + \eta_r^-)\eta_a = \eta_r^+ \left(\frac{\tilde{\rho}_p}{\tilde{\rho}_c^+}\right)^{H_a} + \eta_r^- \left(\frac{\tilde{\rho}_p}{\tilde{\rho}_c^-}\right)^{H_a}. \tag{S16}$$

Therefore, the precise position of the dynamical equilibrium point $(\eta_a,\ \tilde{\rho}_p)$ can be determined from $\tilde{\rho}_p \eta_a = 1$ and Eq. (S16). In particular, the dynamic equilibrium point of a 500 mM oscillatory stimulus is equivalent to the equilibrium point of a 222 mM constant hyperosmotic shock. Surprisingly, the entire trajectories in the internal state space for an oscillatory stimulus and its equivalent hyperosmotic shock almost coincide (Figure S2E). Indeed, the protein density and growth rate followed by the equivalent constant hyperosmotic shock are essentially the time-averaged results of the oscillatory one (Figure S2F).

For walled cells under oscillatory perturbation, due to the extra dynamics of turgor pressure, the target equilibrium point $(\tilde{\rho}_c^{H_a/(H_a+1)}, \tilde{\rho}_c^{-H_a/(H_a+1)})$ moves along the curve $\tilde{\rho}_p \eta_a = 1$ (Movie S4 and Figure S6). The steady state in the internal state space is still a periodic circle around a dynamical equilibrium point on the curve $\tilde{\rho}_p\eta_a = 1$, similar to the case of unwalled cells.

#### E. Cell-wall synthesis regulation is necessary for supergrowth

In this section, we discuss the conditions of the supergrowth phenomenon. Specifically, we focus on the scenario where the cell has undergone sufficient oscillatory osmotic cycles before switching back to the reference growth medium. Our conclusions are also valid for a constant hypoosmotic shock. By definition, the overall growth rate of the entire cytoplasmic volume is the weighted average of the dry-mass growth rate and the free-volume volume growth rate $\mu = (1-f)\mu_r + f\mu_f$. We remark that the condition of supergrowth can be well approximated by the instantaneous condition: $\mu_f > \mu_r$ because $\mu_r$ only depends on the protein density, which changes much less significantly than the free-volume growth rate (Figure S9).

We rewrite the dynamics of the internal osmotic pressure, Eq. (18c) in the main text, in terms of the internal state $(\tilde{\rho}_p, \eta_a)$:

$$\dot{\Pi}_{in} = (\eta_a\tilde{\rho}_p\mu_r - \mu_f)\Pi_{in} \tag{S17}$$

We first discuss the case of an unwalled cell. After sufficient cycles of osmotic oscillation, the internal state $(\tilde{\rho}_p, \eta_a)$ converges to a periodic steady state that lies on $\tilde{\rho}_p\eta_a = 1$ (Figure S2E). After removing the oscillatory stimulus, the internal state leaves the curve $\tilde{\rho}_p\eta_a = 1$ and returns to this curve after fully adapting to the reference growth media. During this adaptation process, the internal osmotic pressure constantly equals the external one because the turgor pressure is zero. Therefore, $\dot{\Pi}_{in} = 0$ in Eq. (S17), the free-volume growth rate takes the form of

$$\mu_f = \mu_r\eta_a\tilde{\rho}_p. \tag{S18}$$

We note that the internal state $(\tilde{\rho}_p, \eta_a)$ always follows a counterclockwise trajectory (Figure S2A, C). Therefore, the unwalled cell inevitably enters the region of $\tilde{\rho}_p\eta_a < 1$ after removing the oscillatory stimulus (Figure S3A), and the supergrowth phenomenon cannot occur according to Eq. (S18) (the upper panel of Figure S3B).

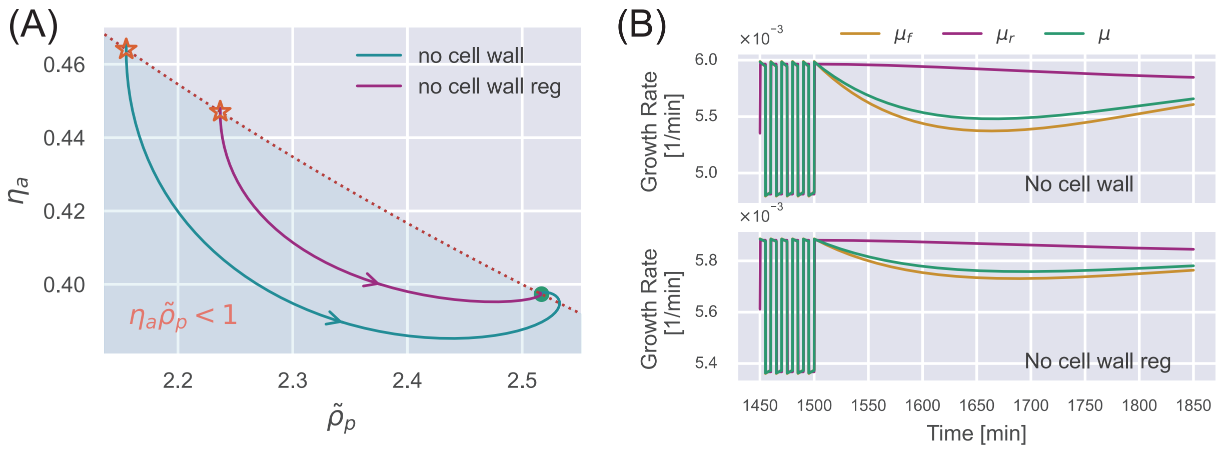


FIG. S3. (A) An oscillation stimulus is initially applied in the simulation until the cell reaches the periodic steady state (stars). The external osmolarity is then switched to the reference growth medium. The internal state evolves towards the steady state in the reference growth medium. Here, we show the results of an unwalled cell (blue curve) and a walled cell deficient in cell-wall synthesis regulation (purple curve). (B) The time dependence of the various growth rates corresponding to the simulations in (A).

Next, we discuss a walled cell deficient in cell-wall synthesis regulation. In this case, the growth rate of the relaxed cell-wall volume is always equal to the growth rate of dry mass, $\mu_{cw} = \mu_r$. According to Eq. (18d) in the main text, the time dependence of turgor pressure is given by

$$\dot{\sigma} = G\dot{\epsilon} = f(G+\sigma)(\mu_f - \mu_r). \tag{S19}$$

To maintain osmotic balance during the adaptation period, $\dot{\Pi}_{in} = \dot{\sigma}$, the free-volume growth rate must satisfy

$$\mu_f = \frac{f(G+\sigma) + \eta_a \tilde{\rho}_p \Pi_{in}}{f(G+\sigma) + \Pi_{in}} \mu_r. \tag{S20}$$

For the same reason as an unwalled cell, the internal state of a walled cell deficient in cell-wall synthesis regulation enters the region of $\tilde{\rho}_p \eta_a < 1$ after removing the oscillatory stimulus (Figure S3A). According to Eq. (S20), there is no supergrowth phenomenon after the oscillation stimulus (the lower panel of Figure S3B). In summary, cell-wall synthesis regulation is necessary for supergrowth.

### F. The growth rate peak during the supergrowth phase

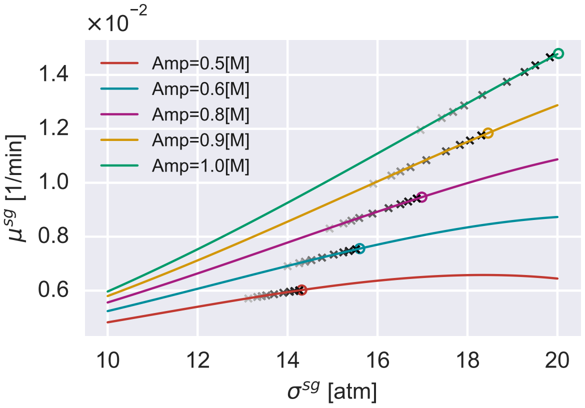


FIG. S4. The relationship between the growth rate peak $\mu^{sg}$ and the turgor pressure at the growth rate peak $\sigma^{sg}$ under different amplitudes of hypoosmotic shock. The solid lines are the theoretical predictions, Eq. (S23). The open circles are obtained by approximating the turgor pressure as the immediate value after the shock $\sigma^f$ in Eq. (S23). The gray crosses represent the peak growth rates obtained by direct simulations using different $\tau^+_{cw}$. As the color goes from light to dark, $\tau^+_{cw}$ gradually decreases to zero.

Here, we consider the case of a constant hypoosmotic shock and derive an analytical expression of the growth rate peak $\mu^{sg}$. As discussed in the main text, the abrupt water influx right after the shock rapidly stretches the cell wall, resulting in an elevated turgor pressure. One should note that the ratio of the dry-mass growth rate before and after the shock is equal to that of the relaxed cell-wall volume, both set by the crowding effect,

$$\frac{\mu^f_{cw}}{\mu^i_{cw}} = \frac{\mu^f_r}{\mu^i_r} = \frac{\eta^f_r}{\eta^i_r}. \tag{S21}$$

Meanwhile, the timescale $\tau^+_{cw}$ governing the up-regulation of cell-wall synthesis efficiency $\eta_{cw}$ to its target value $(\sigma/\sigma_c)^{H_{cw}}$ is considerably smaller than the timescale associated with the osmoregulation process. Consequently, during the timescale of supergrowth, cells can be considered effectively deficient in osmoregulation, producing osmolyte molecules and dry mass at the same rate. Thus, we can simplify the time dependence of the internal osmotic pressure as $\dot{\Pi}_{in} = (\mu_r - \mu_f)\Pi_{in}$. Meanwhile, the water flux is physically constrained to ensure osmotic balance: $\dot{\Pi}_{in} = \dot{\sigma}$, leading to the expression:

$$\mu_f = \mu_r + \frac{1}{f + \frac{\Pi_{in}}{G+\sigma}}(\mu_{cw} - \mu_r). \tag{S22}$$

We point out that the timescales of the relaxation dynamics of $f$, $\Pi_{in}$, and $\sigma$ are all set by the doubling time (Figure S10); therefore, we can approximate them as constant for a short timescale. When the cell-wall synthesis efficiency $\eta_{cw} = \mu_{cw}/\mu_r$ reaches its target value, which is also its maximum value (Figure 3B in the main text), the ratio $\mu_f/\mu_r$ reaches its maximum value according to Eq. (S22), giving rise to the growth rate peak $\mu^{sg}$. Substituting $\mu_{cw} = \mu_r(\sigma/\sigma_c)^{H_{cw}}$ into Eq. (S22) and employing the definition $\mu = f\mu_f + (1-f)\mu_r$, we obtain the expression of $\mu_{sg}$:

$$\mu^{sg} = \mu_r \left\{ 1 + \frac{f}{f + \frac{\Pi_{in}}{\sigma+G}} \times \left[ \left(\frac{\sigma}{\sigma_c}\right)^{H_{cw}} - 1 \right] \right\}. \tag{S23}$$

All the variables on the right side of Eq. (S23) are calculated at the growth rate peak.

Owing to the gradual restoration of turgor pressure, the turgor pressure at the growth rate peak differs from the immediate value after the hypoosmotic shock. Through direct simulations, we acquire the precise value of $\sigma$ at the growth rate peak. Given the value of $\sigma$, the corresponding values of $\Pi_{in}$, $\mu_r$, and $f$ can be inferred. During the adaptation period, osmotic balance is maintained such that $\Pi_{in} = \Pi^f_{out} + \sigma$. Given $\Pi_{in}$, one can infer $\rho_p$ through Eq. (S10) since osmoregulation is effectively inactive, from which we calculate the $\eta_r$ factor and the dry-mass growth rate $\mu_r$. $f$ is subsequently derived from Eq. (S6). Consequently, $\sigma$ is the only degree of freedom on the right-hand side of Eq. (S23). We compare this analytical expression with direct simulations of *S. pombe* at different up-regulation timescales of the cell-wall synthesis process (Figure S4). As $\tau^+_{cw}$ tends to zero, the growth rate peak can be well approximated by the immediate value after the hypoosmotic shock $\sigma^f$. This extreme case is a good approximation for wild-type cells, since the timescale of the turgor pressure recovery process is set by the doubling time (Figure S10), much longer than $\tau^+_{cw}$. Indeed, a strong correlation exists between the growth rate peak and the turgor pressure right after the shock (Figure 3E of the main text).

#### G. The overshoot of turgor pressure after a single oscillation

During a single osmotic oscillation, the external osmolarity shifts to a hyperosmotic environment and then switches back to the reference growth medium (upper panel of Figure S5). We represent the turgor pressure jump resulting from the first osmolarity shift as $\sigma^{f,1} - \sigma^{i,1}$ and the corresponding jump from the second shift as $\sigma^{f,2} - \sigma^{i,2}$ (Figure S5). In Figure 4E of the main text, we show that the turgor pressure overshoot after the single oscillation $\sigma^{f,2} - \sigma_c$ can be approximated by the turgor recovery $\delta\sigma$ during the hyperosmotic period.

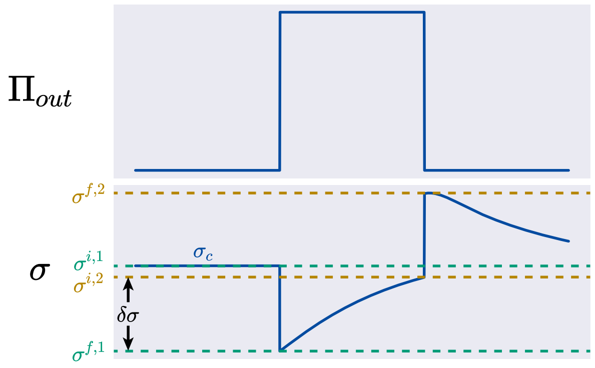


FIG. S5. The upper panel shows the time dependence of the external osmolarity, and the lower panel shows the trajectory of the turgor pressure under a mild osmotic oscillation.

To prove the statement $\delta\sigma \approx \sigma^{f,2} - \sigma_c$, we only need to prove that the absolute magnitudes of the two turgor pressure jumps are nearly identical (Figure S5). The analytical expression for the turgor pressure after the jump is shown in Eq. (S4), where $\sigma^f$ is only related to $\sigma^i$ and $\epsilon^i$ before the shock. For a mild oscillation amplitude, the physiological properties of the cell are not much perturbed. In particular, the turgor pressure has almost recovered to the steady-state value $\sigma_c$ (Figure S5). We denote $\Delta\sigma = \sigma^f - \sigma^i$ and find that for a mild osmotic perturbation such that $\Delta\Pi_{in} = \Delta\sigma + \Delta\Pi_{out} \ll \Pi^i_{in} = \sigma^i + \Pi^i_{out}$, Eq. (S4) can be simplified as:

$$\frac{\Delta\sigma}{\sigma^i} = -\left(1 + G/\sigma^i\right) f^i \frac{\Delta\Pi_{out} + \Delta\sigma}{\Pi^i_{in}}. \tag{S24}$$

It is evident that $\Delta\sigma$ is directly proportional to $\Delta\Pi_{out}$. Given that $\Delta\Pi^2_{out} = -\Delta\Pi^1_{out}$, the turgor pressure jumps cancel each other out for a mild osmotic oscillation.

[1] G. Lesage and H. Bussey, Cell wall assembly in saccharomyces cerevisiae, Microbiology and molecular biology reviews **70**, 317 (2006).

[2] F. M. Klis, P. Mol, K. Hellingwerf, and S. Brul, Dynamics of cell wall structure in saccharomyces cerevisiae, FEMS microbiology reviews **26**, 239 (2002).

[3] S. Cayley, B. A. Lewis, H. J. Guttman, and M. T. Record Jr, Characterization of the cytoplasm of escherichia coli k-12 as a function of external osmolarity: implications for protein-dna interactions in vivo, Journal of molecular biology **222**, 281 (1991).

[4] Y. T. Chong, J. L. Koh, H. Friesen, S. K. Duffy, M. J. Cox, A. Moses, J. Moffat, C. Boone, and B. J. Andrews, Yeast proteome dynamics from single cell imaging and automated analysis, Cell **161**, 1413 (2015).

[5] E. A. Yamada and V. C. Sgarbieri, Yeast (saccharomyces cerevisiae) protein concentrate: preparation, chemical composition, and nutritional and functional properties, Journal of agricultural and food chemistry **53**, 3931 (2005).

[6] A. Miermont, F. Waharte, S. Hu, M. N. McClean, S. Bottani, S. Léon, and P. Hersen, Severe osmotic compression triggers a slowdown of intracellular signaling, which can be explained by molecular crowding, Proceedings of the National Academy of Sciences **110**, 5725 (2013).

[7] A. T. Molines, J. Lemière, M. Gazzola, I. E. Steinmark, C. H. Edrington, C.-T. Hsu, P. Real-Calderon, K. Suhling, G. Goshima, L. J. Holt, et al., Physical properties of the cytoplasm modulate the rates of microtubule polymerization and depolymerization, Developmental Cell **57**, 466 (2022).

[8] E. Zhou, X. Trepat, C. Park, G. Lenormand, M. Oliver, S. Mijailovich, C. Hardin, D. Weitz, J. Butler, and J. Fredberg, Universal behavior of the osmotically compressed cell and its analogy to the colloidal glass transition, Proceedings of the National Academy of Sciences **106**, 10632 (2009).

[9] H. Ting-Beall, D. Needham, and R. Hochmuth, Volume and osmotic properties of human neutrophils, Blood **81**, 2774 (1993).

[10] J. Lemière and F. Chang, Quantifying turgor pressure in budding and fission yeasts based upon osmotic properties, bioRxiv 10.1101/2023.06.07.544129 (2023).

[11] E. Atilgan, V. Magidson, A. Khodjakov, and F. Chang, Morphogenesis of the fission yeast cell through cell wall expansion, Current Biology **25**, 2150 (2015).

[12] D. S. Cayley, H. J. Guttman, and M. T. Record, Biophysical characterization of changes in amounts and activity of escherichia coli cell and compartment water and turgor pressure in response to osmotic stress, Biophysical journal **78**, 1748 (2000).

[13] B. D. Knapp, P. Odermatt, E. R. Rojas, W. Cheng, X. He, K. C. Huang, and F. Chang, Decoupling of rates of protein synthesis from cell expansion leads to supergrowth, Cell systems **9**, 434 (2019).

Movie S1: The trajectory in the internal state space for an wide-type cell during a 500 mM hyperosmotic shock. The movie shown here corresponds to the dynamics depicted in Figure 3C of the main text.

Movie S2: The trajectory in the internal state space for an wide-type cell during a 500 mM hypoosmotic shock. The movie shown here corresponds to the dynamics depicted in Figure 3D of the main text.

Movie S3: The trajectory in the internal state space for an unwalled cell during a 500 mM osmotic oscillation with a 10-minute period. The movie shown here corresponds to the dynamics depicted in Figure S2E.

Movie S4: The trajectory in the internal state space for an wide-type cell during a 500 mM osmotic oscillation with a 10-minute period. The movie shown here corresponds to the dynamics depicted in Figure S6.

| Symbol | Description |
|---|---|
| $V$ | total cytoplasmic volume |
| $V_f$ | free volume: cytoplasmic volume occupied by free water |
| $V_b$ | bound volume: cytoplasmic volume occupied by dry mass and bound water |
| $V_{bw}$ | volume occupied by bound water |
| $V_{bd}$ | dry volume: volume occupied by dry mass |
| $V_{cw}$ | relaxed cell wall volume |
| $f$ | free volume fraction |
| $\alpha$ | bound volume per total protein mass |
| $\Pi_{in}$ | cytoplasmic osmotic pressure |
| $\Pi_{out}$ | external osmotic pressure |
| $\Pi_{in,c}$ | critical cytoplasmic osmolarity where cell growth arrests |
| $\Pi_{out,c}$ | critical external osmolarity where cell growth arrests |
| $\sigma$ | turgor pressure |
| $G$ | cell wall elastic modulus |
| $\epsilon$ | elastic strain of the cell wall |
| $k_w$ | water permeability of the cell membrane |
| $N_a$ | number of osmolyte molecules in cytoplasm |
| $m_p$ | total mass of the proteome |
| $m_{p,a}$ | mass of the osmolyte-producing protein |
| $m_{p,r}$ | mass of the ribosomal protein |
| $\rho_p$ | protein density |
| $\rho_c$ | critical protein density where cell growth arrests |
| $\tilde{\rho}_p$ | normalized protein density |
| $\tilde{\rho}_c$ | normalized critical protein density |
| $\bar{\rho}_p$ | normalization factor in protein density |
| $\phi_a$ | mass fraction of the osmolyte-producing protein in total proteome |
| $\phi_r$ | mass fraction of the ribosomal protein in total proteome |
| $\chi_a$ | fraction of ribosome translating osmolyte-producing protein |
| $\chi_r$ | fraction of ribosome translating ribosomal protein |
| $\chi_a^{\max}$ | largest possible fraction of ribosome translating osmolyte-producing protein |
| $H_a$ | sensitivity of osmoregulation to intracellular crowding |
| $H_{cw}$ | sensitivity of cell-wall synthesis regulation to turgor pressure |
| $\tau_{cw}^{\pm}$ | timescale of up(down)-regulation of cell-wall synthesis regulation |
| $k_a$ | osmolyte production rate |
| $k_a^{\max}$ | maximum osmolyte production rate |
| $k_r$ | ribosomal protein production rate |
| $k_r^{\max}$ | maximum ribosomal protein production rate |
| $\eta_a$ | efficiency of osmoregulation |
| $\eta_{cw}$ | efficiency of cell-wall synthesis regulation |
| $\eta_r$ | crowding factor |
| $\mu$ | growth rate of total volume |
| $\mu_r$ | growth rate of ribosomal protein (dry mass) |
| $\mu_f$ | growth rate of free volume |
| $\mu_{cw}$ | growth rate of relaxed cell wall volume |
| $\mu^{sg}$ | peak growth rate of total volume during supergrowth phase |

TABLE S2. A summary of the symbols involved in our model.

| Parameters | S. pombe (WT) | S. pombe (no osmoregulation) | S. pombe (no CW synthesis regulation) | S. pombe (no CW) |
|---|---|---|---|---|
| $\Pi_{out}^*$ | 0.2 [Osm] | 0.2 [Osm] | 0.2 [Osm] | 0.2 [Osm] + 10 [atm] |
| $\sigma^*$ | 10 [atm] | 10 [atm] | 10 [atm] | |
| $\rho_p^*$ | 0.104 [g/ml] | 0.104 [g/ml] | 0.104 [g/ml] | 0.104 [g/ml] |
| $\mu^*$ | 0.35 [1/h] | 0.35 [1/h] | 0.35 [1/h] | 0.35 [1/h] |
| $f^*$ | 0.788 | 0.788 | 0.788 | 0.788 |
| $\epsilon^*$ | 0.584 | 0.584 | 0.584 | |
| $\alpha$ | 2.60 [ml/g] | 2.60 [ml/g] | 2.60 [ml/g] | 2.60 [ml/g] |
| $G$ | 17.1 [atm] | 17.1 [atm] | 17.1 [atm] | |
| $\Pi_{out,c}$ | 3.5 [Osm] | 1.15 [Osm] | 3.5 [Osm] + 10 [atm] | 3.5 [Osm] + 10 [atm] |
| $k_w$ | 100 [1/(min atm)] | 100 [1/(min atm)] | 100 [1/(min atm)] | 100 [1/(min atm)] |
| $\rho_c$ | 0.267 [g/ml] | 0.267 [g/ml] | 0.267 [g/ml] | 0.267 [g/ml] |
| $\sigma_c$ | 10 [atm] | 10 [atm] | 10 [atm] | |
| $H_r$ | 3.03 | 3.03 | 3.03 | 3.03 |
| $H_a$ | 0.974 | 0 | 0.974 | 0.974 |
| $k_r^{\max}\chi_r$ | 0.371 [1/h] | 0.371 [1/h] | 0.371 [1/h] | 0.371 [1/h] |
| $k_B T k_a^{\max}\chi_a^{\max}$ | 2.25 [(atm ml)/(g min)] | 0.894 [(atm ml)/(g min)] | 2.25 [(atm ml)/(g min)] | 2.25 [(atm ml)/(g min)] |
| $\tau_{cw}^-$ | 0.1 [min] | 0.1 [min] | | |
| $\tau_{cw}^+$ | 12.5 [min] | 12.5 [min] | | |
| $H_{cw}$ | 1.7 | 1.7 | 0 | |

TABLE S3. Comparison of the model parameters between defective cells and intact cells. Variables with $*$ are for the reference growth medium.

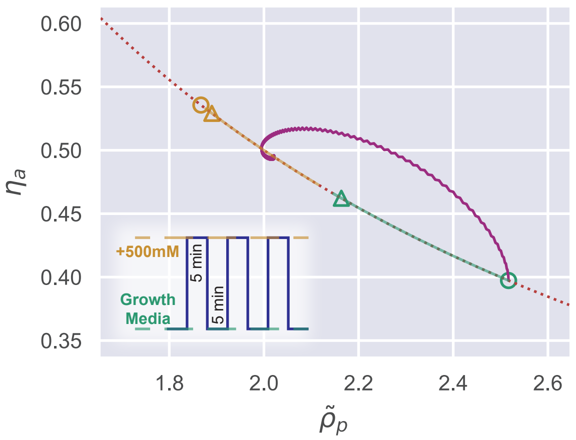


FIG. S6. Trajectory of a wild-type walled cell in the internal state space during a 500 mM osmotic oscillation. During the oscillation, the equilibrium points move along the dotted curve $\tilde{\rho}_p \eta_a = 1$. The circles indicate the equilibrium points during the first oscillation cycle, while the triangles indicate their positions after a large enough number of oscillations.

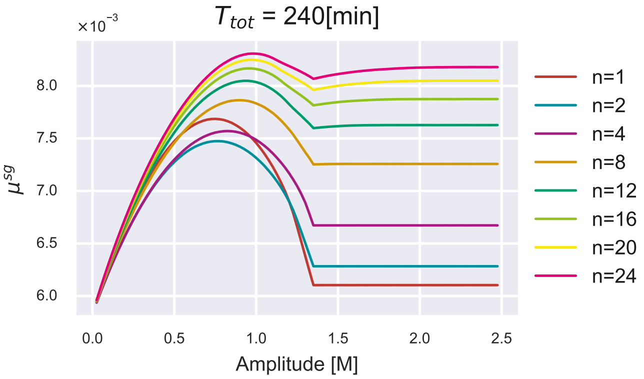


FIG. S7. Growth rate peak $\mu^{sg}$ *vs.* oscillation amplitude under different numbers of oscillation period. The total duration of the oscillation stimulus is fixed.

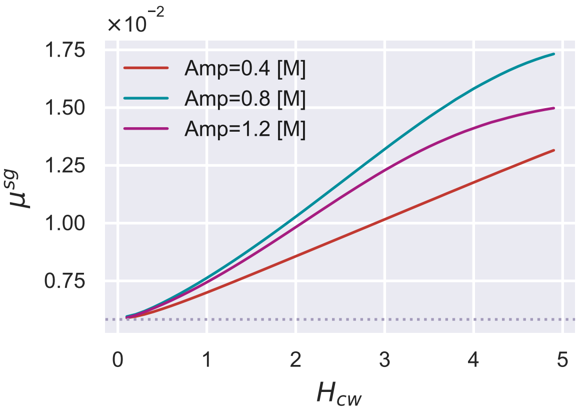


FIG. S8. Given 24 cycles of osmotic oscillations with a period of 10 minutes, we vary the parameter $H_{cw}$, which dictates the sensitivity of cell-wall synthesis regulation. A higher growth rate peak $\mu^{sg}$ is observed as the sensitivity $H_{cw}$ increases, irrespective of the oscillation amplitude. The dotted line represents the growth rate in the reference growth medium.

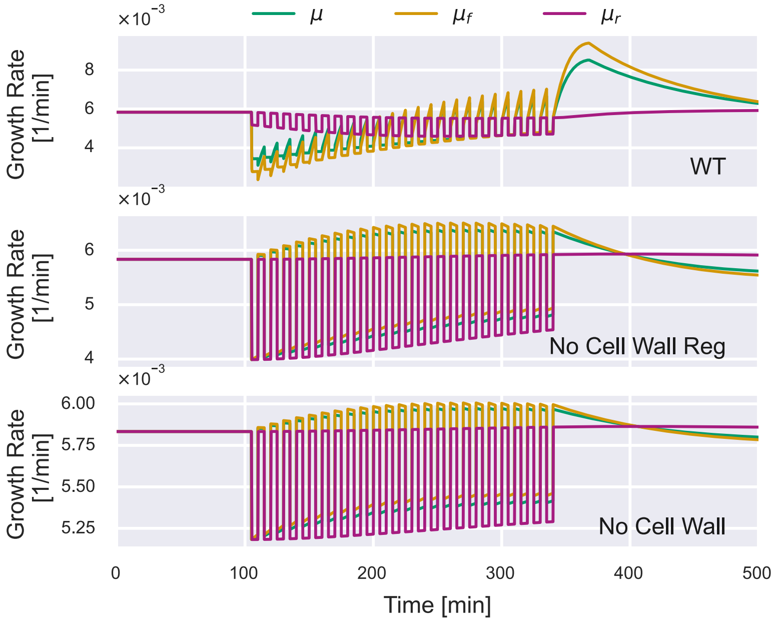


FIG. S9. Time dependence of the free-volume growth rate $\mu_f$, the dry-mass growth rate $\mu_r$, and the overall growth rate $\mu$ under 500 mM osmotic oscillations. Here, we show the results of an intact WT cell (upper panel), a walled cell deficient in cell-wall synthesis regulation, and an unwalled cell.

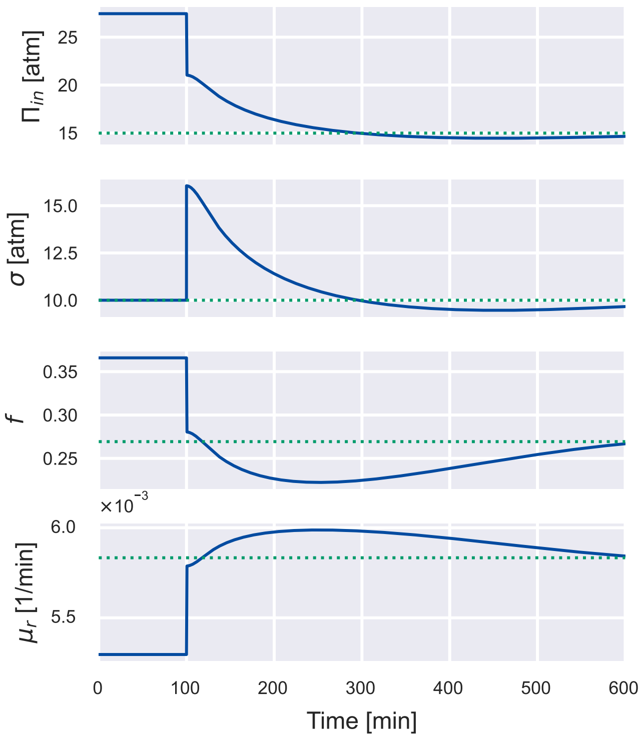


FIG. S10. Simulation of a cell undergoing a constant 500 mM hypoosmotic shock, the same simulation as Figure 3B in the main text. Here, we show the time dependence of the internal osmotic pressure $\Pi_{in}$, the turgor pressure $\sigma$, the free volume fraction $f$, and the dry-mass growth rate $\mu_r$. All these quantities relax to their steady-state values (dotted lines) on the timescale set by the doubling time.

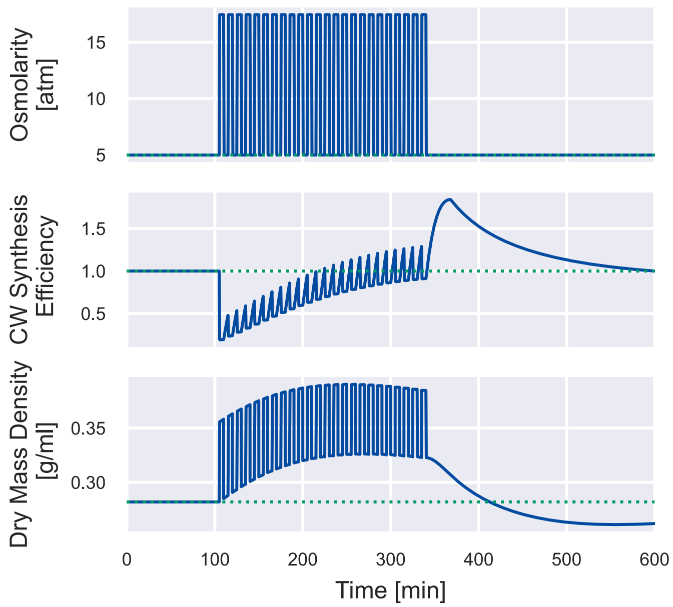


FIG. S11. Simulation of a wild-type cell undergoing 24 cycles of 500 mM osmotic oscillation with a 10-min period, the same simulation as Figure 4A in the main text. We plot the cell-wall synthesis efficiency $\eta_{cw}$ and the dry-mass density $\hat{\rho}_d$ for better comparison with experimental data [13].
